# Endonucleosis mediates internalization of cytoplasm into the nucleus in senescent cells

**DOI:** 10.1101/2023.11.12.566736

**Authors:** Ourania Galanopoulou, Evangelia C. Tachmatzidi, Elena Deligianni, Dimitris Botskaris, Kostas C. Nikolaou, Sofia Gargani, Yannis Dalezios, Georges Chalepakis, Iannis Talianidis

## Abstract

Cellular senescence is driven by diverse effector programs, leading to irreversible growth arrest, DNA damage and complex secretomes. Here we show that, in liver-specific Setd8-KO mice, after mitogen treatment, a significant number of hepatocytes become senescent and display unusual features such as enlarged nuclei, chromosomal hyperploidy and nuclear engulfments progressing to the formation of intranuclear vesicles. These vesicles contain glycogen, cytoplasmic proteins and even entire organelles. We term this process “endonucleosis”. Experiments with Setd8/Atg5 double knockout mice, demonstrated that endonucleosis requires the function of the autophagy machinery. Endonucleosis and hyperploidization are temporary, early features of senescence. Larger vesicles brake down into microvesicles over time and are eventually eliminated. The results reveal a senescence phenotype, which function as part of survival mechanisms to prevent necrotic death.

## Main Text

Setd8 is the sole enzyme catalyzing H4K20 monomethylation (*1, 2*). H4K20Me_1_ is used as substrate by Suv4-20h1/2 to generate H4K20Me_2_ and H4K20Me_3_, which facilitate nucleosomal folding and heterochromatin formation (*2*). Previous studies have established the pivotal role of Setd8 in the regulation of genome integrity (*3–6*), mitotic chromatin condensation (*7, 8*), replication licensing (*9, 10*) and DNA damage response (*2–6*). We have previously shown that hepatocyte-specific inactivation of Setd8, leads to cell division-dependent extensive DNA damage and massive necrotic cell death (*11*). As the majority of hepatocytes in the adult liver are in the G0 phase, necrotic cell death in 2-4 months old mice is initially observed in microscopic focal areas of the liver. The initial cell death triggers a local regenerative process, whereby neighboring cells enter the cell cycle and accelerate the death of the existing Setd8-deficient hepatocytes, resulting in the expansion of the necrotic zone. Within the necrotic areas of the liver, we observed enlarged hepatocytes and accumulation of inflammatory cells (*11*). Large hepatocytes appeared homogenously in all areas of the liver after either partial hepatectomy, which induces cell cycle entry (*11*), or under metabolic stress conditions, such as fasting, high fat diet or pyruvate challenge (*12*). At 8 to 12 months of age, Setd8-deficient mice develop full-blown spontaneous liver tumors comprising transformed hepatocytes with cancer stem cell properties (*11–13*). The apparent discrepancy between Setd8 deficiency induced necrosis and the late onset hepatocellular carcinoma prompted us to further investigate and characterize the process of hepatocyte enlargement triggered by spontaneous or induced cell proliferation.

### Internalization of cytoplasm into the nuclei by endonucleosis

Treatment of liver-specific Setd8 knockout mice (Setd8-LKO) with TCPOBOP, a widely used CAR agonist that induces robust hepatocyte proliferation (*14–16*), resulted in synchronous and homogeneous enlargement of hepatocytes within 24 hours and an approximately 8-fold increase in the nuclear size (Fig. 1A and fig. S1A to G). Enlarged hepatocytes occupied almost the entire liver space along with a significant enrichment of small mononuclear cells (Fig. 1A and fig. S1E and G). Indicative of the massive cell death in Setd8-LKO livers, only about 10% of the hepatocyte population remained in Setd8-LKO livers compared to wild-type mice (fig. S1D). We examined hundreds of high-power fields in sections within the 24-hour period but could not detect any nuclear or cellular configuration that would suggest cell or nuclear fusion. Therefore, we anticipate that the surviving cells and nuclei expanded physically.

**Fig. 1.**
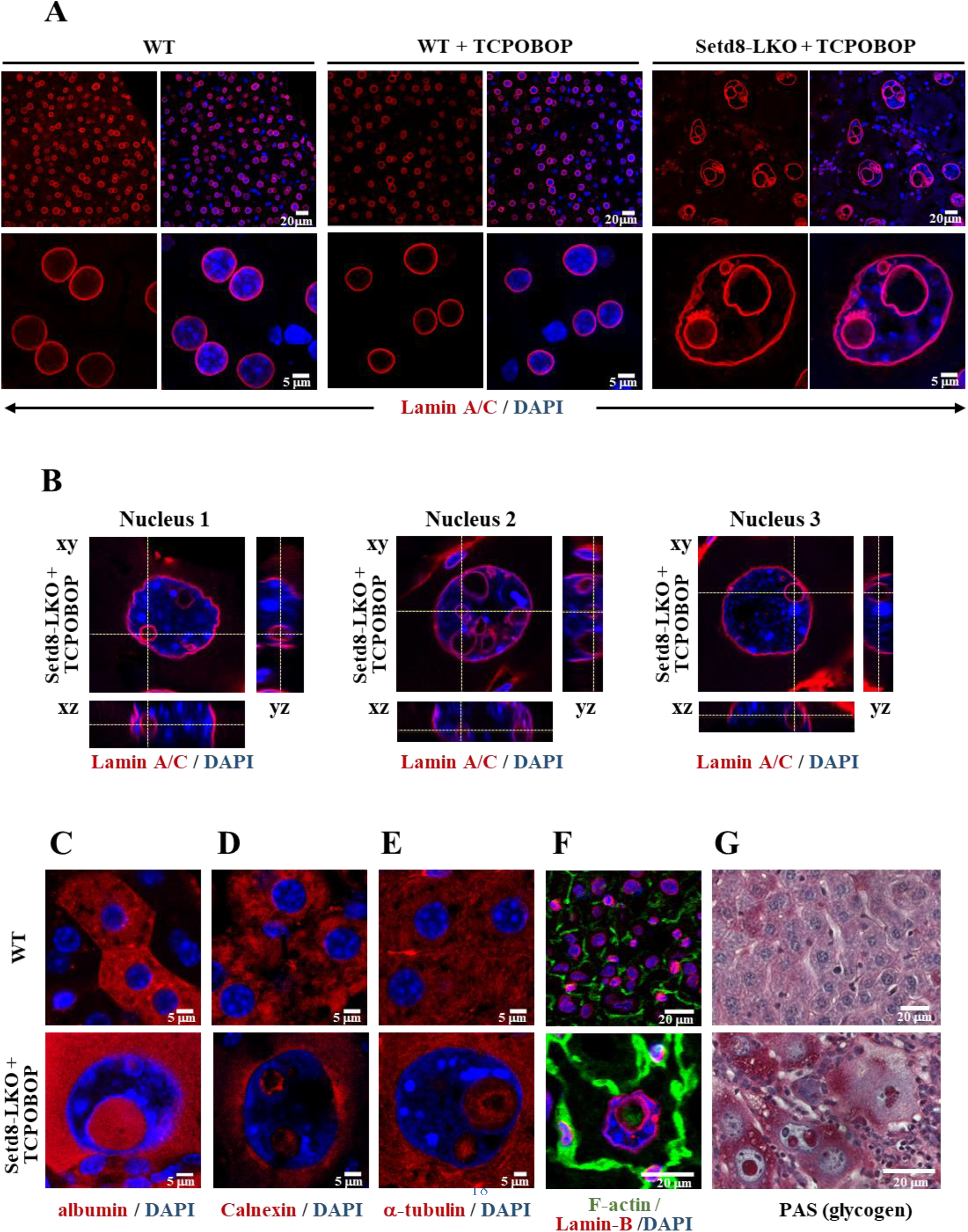
Intranuclear lamin-coated vesicles containing cytoplasmic proteins in Setd8-LKO hepatocytes. (**A**) LaminA/C antibody immunostaining of liver sections from wild type (WT) mice and from wild type or Setd8^lox/lox^/AlbCre (Setd8-LKO) mice 24 hours after treatment with TCPOBOP (WT+TCPOBOP and Setd8-LKO+TCPOBOP). Fluorescent images of the cells are shown at different magnifications. (**B**) Orthogonal views of confocal z-stacks showing 3D morphology of internal vesicles in 3 typical Lamin A/C-stained nuclei of TCPOBOP-treated Setd8-LKO hepatocytes. (**C-F**), Representative immunofluorescence images with antibodies against albumin (**C**), calnexin (**D**), a-tubulin (**E**) and F-actin (**F**). (**G**) Representative image of periodic acid-Schiff (PAS) staining of glycogen.

Most of the enlarged nuclei contained Lamin A/C and Lamin B1-coated heterogeneously-sized vesicles (Fig. 1A and Fig. S1A). The presence of Sun2, a core component of the inner nuclear membrane-associated LINC complex, that connects the nuclear lamina to actin cytoskeleton (*17*), further confirmed that the membranes coating the vesicles originated from the nuclear envelope (fig. S1B). Lamin A/C polymers were thinner but had proportionally higher staining intensity in Setd8-LKO cells than in wild-type cells, as revealed by measuring the thickness and the staining intensity of the lamin A/C signal in a multitude of regions (fig. S2A and B). These data point to a high level of elasticity of the nuclear lamina with stretched and denser lamin polymers in Setd8-LKO hepatocytes. Although the lamin A/C protein content may not differ locally, its overall levels per individual cell were greatly increased in Setd8-LKO hepatocytes, considering the greatly increased nuclear size and the presence of the numerous intranuclear lamin-coated vesicles. 3D reconstruction of confocal sections demonstrated that the vesicles were positioned entirely within the nuclear space (Fig.1B, and fig. S2C). The intranuclear vesicles contained cytoplasmic proteins, such as albumin, calnexin, α-tubulin, F-actin filaments and glycogen (Fig. 1C to G), but excluded nuclear proteins, such as HNF4α (fig. S1B) and DNA (Fig. 1A and B).

Transmission electron microscopy analyses captured several steps of the cytoplasm internalization process. We observed micro-extensions of the nuclear membrane, membrane invaginations towards the nuclear interior and cytoplasm internalization via fusion of the inwardly folded nuclear membrane. Fusion occurred at the tip of the engulfed areas, thus sequestering cytoplasmic materials into intranuclear vesicles (Fig. 2A and B, and fig. S3A). As a result of membrane folding towards the nuclear interior, lamina-associated chromatin is positioned at the external side of the resulting vesicles (Fig. 2A to C). Moreover, in larger vesicles, we could detect glycogen or even whole cytoplasmic organelles, such as endoplasmic reticulum, mitochondria, peroxisomes and autophagosome structures (Fig. 2C and fig. S3A). We propose to call this process “endonucleosis”, based on its similarity to endocytosis, the process of by which external substances are internalized into the cells (18).

**Fig. 2.**
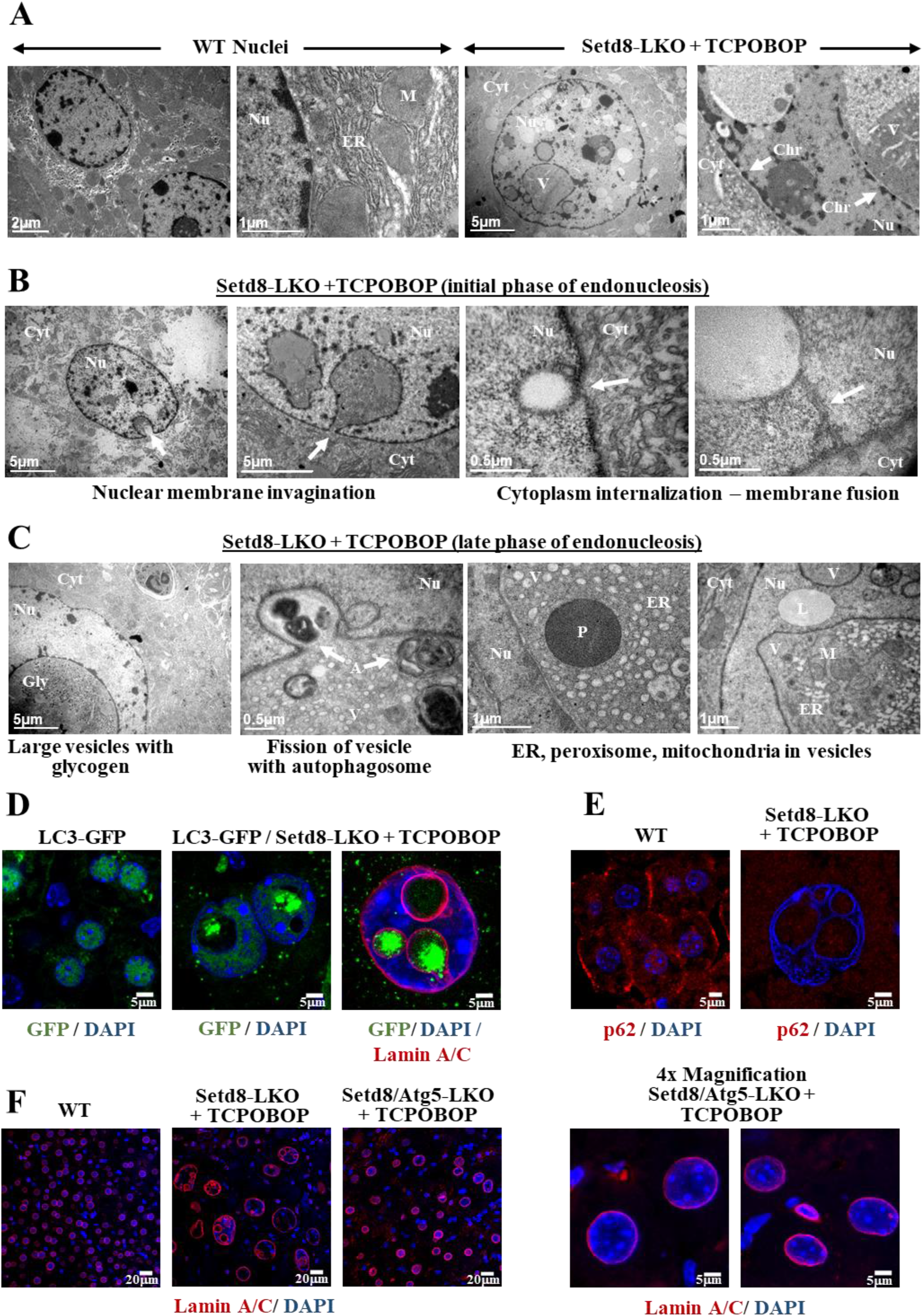
Nuclear membrane folding-mediated internalization of cytoplasmic components and the requirement for autophagy regulator Atg5. **(A-C)** Transmission electron microscopic images of liver sections from wild type and Setd8-LKO mice after 24 hours of TCPOBOP treatment. White arrows indicate: Nu, nucleus; Cyt, cytoplasm; ER, endoplasmic reticulum; M, mitochondria; V, vesicles; Chr, chromatin; Gly, glycogen; A, autophagosome structures; L, lipid droplet. (**D)** Detection of the autophagy marker protein LC3 by GFP fluorescence in LC3-GFP transgenic mice and LC3-GFP/Setd8-LKO mice 24 hours following TCPOBOP treatment. (**E)** Immunofluorescence staining for the autophagy receptor protein p62. (**F)** Lamin A/C staining in wild type, Setd8-LKO and Setd8-LKO/Atg5-LKO double knockout mice after 24 hours of TCPOBOP treatment. Note the lack of intranuclear vesicles and the smaller enlargement of the nuclei in the double knockout mice.

### Endonucleosis requires Atg5 function

We have previously reported that constitutive and persistent activation of AMPK in Setd8-LKO mice leads to the appearance of cytoplasmic autophagic structures in parallel to the rise of the lipidated form of Map1lc3a (LC3) (*11*). Using LC3-GFP transgenic mice (*19*) to monitor LC3 distribution in Setd8-LKO mice, we observed high levels of punctate fluorescence signals within the vesicles, which is characteristic of the activated form of the protein (*20*), as opposed to the homogenously distributed low levels of nuclear LC3 (Fig. 2D). We also observed a parallel decrease in p62 staining signal, which provides additional evidence for elevated autophagy activity in Setd8-LKO mice (Fig. 2E). The punctate form of LC3 was located inside the vesicles. This form of LC3 is involved in autophagic membrane trafficking and the engulfment of cytoplasmic components into autophagosomes. This strongly suggests that endonucleosis may use the activities of the classical autophagy machinery to internalize cytoplasm into the nucleus. This possibility is supported by our findings in TCPOBOP-treated Setd8/Atg5 double knockout mice, where we observed intermediate-sized nuclei, which lacked endonucleotic vesicles (Fig. 2F, fig. S1C and 3B). Atg5 is an indispensable component of both, the canonical and non-canonical autophagy process (*21*). Thus, we conclude that the function of the cytoplasmic autophagy machinery is required for nuclear membrane folding-mediated endonucleosis.

### Hyperploidy and senescence in Setd8-LKO livers

TCPOBOP induces hepatic cell proliferation by promoting the entry into S-phase, of hepatocytes which normally reside at the G0 phase. This results in a transient increase of hepatocyte number and liver size, peaking at 36-48 hours after treatment (*14*). Consistent with previous observations (*14, 15*) in wild-type mice, at the early time point of 24 hours, we observed a slight increase in the number of cells staining positively for the proliferation marker Ki67, without any changes to the average nuclear size and ploidy of the cells (Fig. 3C, and fig. S1C). Surprisingly, the normally diploid or tetraploid hepatocytes became hyperploid in TCPOBOP-treated Setd8-LKO mice and a significant proportion expressed Ki67 (Fig. 3A and C). Moreover, EdU pulse labelling assays detected a number of cells actively replicating DNA (fig. S3C). Propidium iodide (PI) staining assays showed the loss of diploid cells (with 2N DNA content), a reduction in the proportion of tetraploid (4N) cells to 4.5% of the cells and the concomitant appearance of cells with 16N, 32N, 64N, and 128N DNA content, representing ∼95% of the total hepatocyte population (Fig. 3A). Hyperploidy was also confirmed by *in situ* DNA-FISH assays using probes from three different chromosomes (Fig. 3B). The periodicity of the PI staining profile indicates that hyperploidy was generated by consecutive duplication events (endoreduplication) without cell division. This phenomenon could be explained by the role of Setd8 in replication licensing. Previous studies have shown that, in cycling cells, proper recruitment of the origin recognition complex (ORC) to chromatin is mediated by Suv4-2h1/2-catalyzed conversion of H4K20Me_1_ to H4K20Me_2/3_-modified nucleosomes (*9*). It has been postulated that the regulation and timing of H4K20 mono-, di and tri-methylation is critical for proper replication firing, as decreased H4K20Me_1_ and increased H4K20Me_2/3_ levels correlated with aberrant re-replication in Setd8-depleted cells (*10*). Consistent with this possibility, we detected high levels of trimethylated H4K20 by immunostaining in TCPOBOP-treated Setd8-LKO hepatocytes (fig. S3E).

**Fig. 3.**
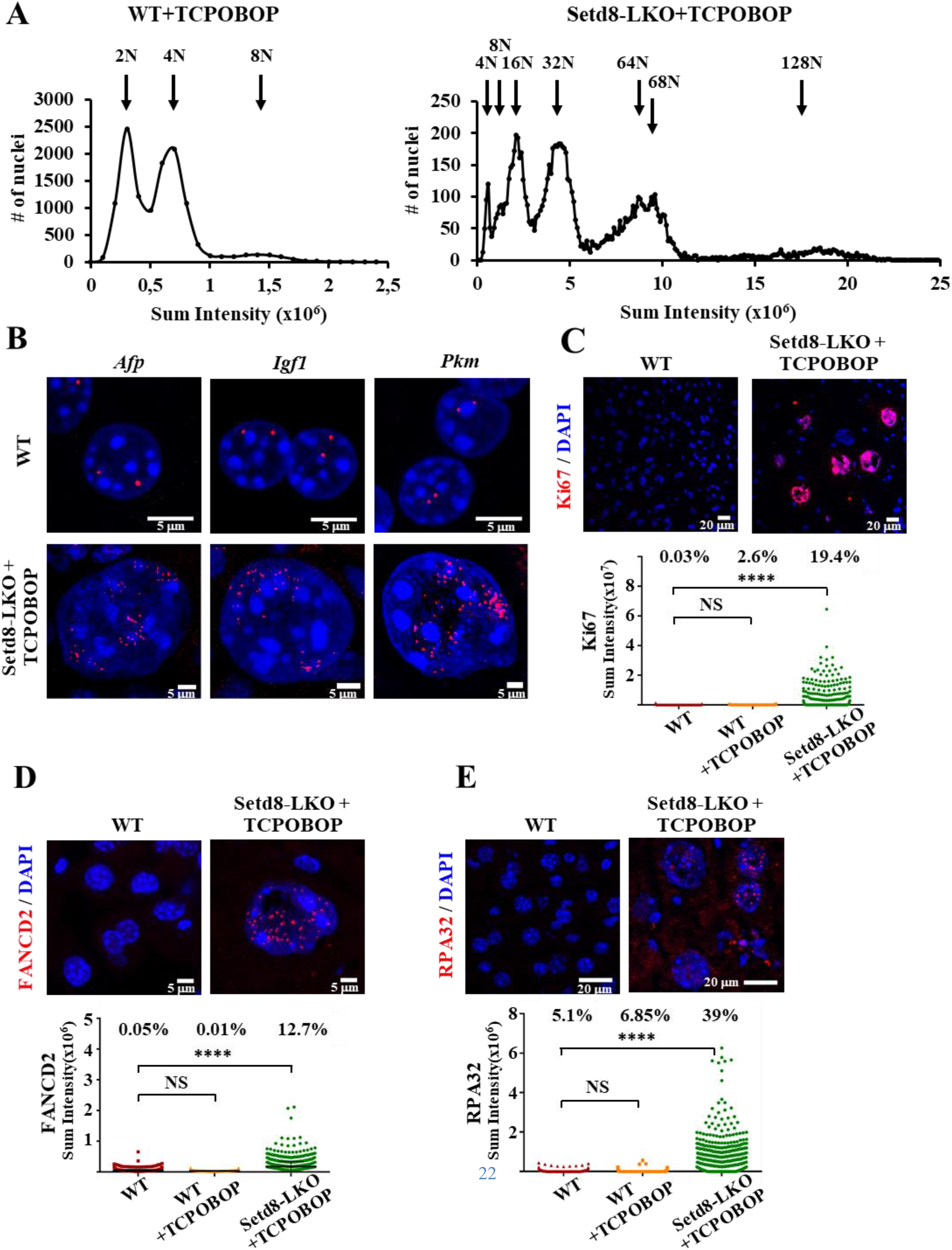
Hyperploidization and replicative stress in Setd8-LKO hepatocytes. (**A**) Quantification of nuclear DNA content in TCPOBOP-treated wild type (n=12217) and Setd8-LKO livers (n=12163) by propidium iodide (PI) staining. Graphs show high-content microscopy (HCM) measurements of the PI staining intensity in individual cells after 24 hours of treatment. Arrows indicate the chromosomal ploidy of the peaks. The percentage of the cells corresponding to the peak areas were as follows: in WT+TCPOBOP, 2N=47.6%; 4N=44.7%, 8N=6.7%; in Setd8-LKO+TCPOBOP, 4N=4.5%; 8N=5.1%, 16N=19.5%; 32N=32.9%; 64N/68N=29.5%, 128N=8.2%. (**B**) Representative DNA FISH images (red) with probes spanning Afp (chr5), Igf1 (chr10) and Pkm2 (chr9) genes. (**C**) Immunofluorescence images of liver sections stained with antibody for the proliferation marker Ki67. The graph shows HCM measurements of the Ki67 staining intensities in individual cells (n=1161 for wild type, n=1226 for WT+TCPOBOP and n=1228 for Setd8-LKO+TCPOBOP). Numbers indicate the percentage of cells with staining intensity above threshold. (**D-E**) Immunofluorescence images of liver sections stained with FANCD2 or RPA32. The graph shows HCM measurements of the staining intensities in individual cells (FANCD2 staining n=2299 for wild type, n=2334 for WT+TCPOBOP and n=2224 for Setd8-LKO+TCPOBOP; RPA32 staining n=3298 for wild type, n=3384 for WT+TCPOBOP and n=2458 for Setd8-LKO+TCPOBOP). Numbers indicate the percentage of cells with staining intensity above threshold. ****p-value<0.0001. NS, Non-significant.

High levels of endoreduplication are expected to be accompanied by replication stress and genome instability. We used FANCD2 and RPA32 immunostaining to monitor replication stress-induced lesions and γH2AX staining for the detection of double-stranded DNA breaks. We could readily detect a significant number of cells with punctate FANCD2, RPA32 and γH2AX staining patterns in Setd8-LKO hepatocytes (Fig. 3D to E and Fig. 4A). These results are consistent with the reported role of Setd8 in the maintaining genome stability (*1–6*).

**Fig. 4.**
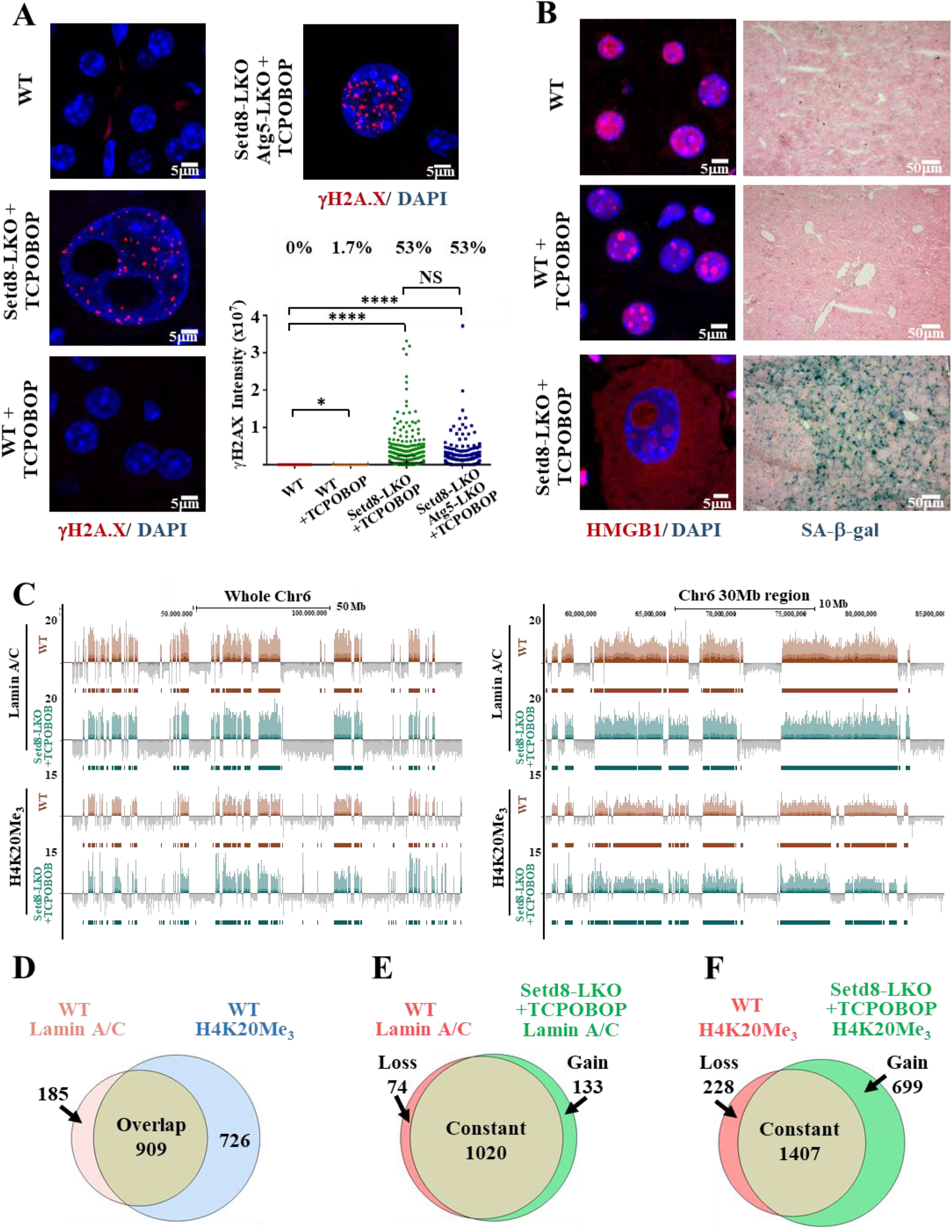
Senescence phenotype, excessive DNA damage and unaltered distribution of lamin-associated genomic domains in Setd8-LKO hepatocytes. (**A**) Representative images of γH2A.X immunostaining of liver sections from wild type and TCPOBOP-treated wild type, Setd8-LKO and Setd8-LKO/Atg5-LKO double knockout mice. The graph shows HCM measurements of the γH2A.X immunofluorescence intensities in individual cells (n=3412 for wild type; n=3382 for Setd8-LKO+TCPOBOP; n=3977 for WT+TCPOBOP and n=3699 for Setd8-LKO/Atg5-LKO+TCPOBOP). Numbers indicate the percentage of cells with staining intensity above threshold. (**B**) Representative images of HMGB1 immunofluorescence and SA-β-galactosidase staining of liver sections 24 hours after TCPOBOP treatment. (**C**) Genome Browser tracks showing normalized CUT&Tag reads of Lamin A/C-associated and H4K20Me_3_-occupied genomic regions on the entire chromosome 6 (left panel) and a 30 Mb region of chromosome 6 (panel at right). (**D**) Venn diagram showing the extent of overlap between Lamin A/C and H4K20Me_3_ CUT&Tag peaks in wild-type liver nuclei. (**E-F**) Venn diagram comparing Lamin A/C (**E**) and H4K20Me_3_ (**F**) CUT&Tag peaks between wild type and Setd8-LKO mice 24 hours after TCPOBOP treatment. “Loss” indicates peaks present only in wild type livers, “Gain” indicates new peaks in Setd8-LKO livers, while “Constant” correspond to peaks found in both, wild type and Setd8-LKO livers. *p-value<0.04; ****p-value<0.0001. NS, Non-significant

The dramatic enlargement of Setd8-LKO hepatocytes, the observed growth arrest, and the extensive DNA damage raised the question of whether these cells escape necrotic death by entering a senescent state (*22, 23*). Strong evidence for the acquisition of a senescent phenotype was provided by the observation of high senescence-associated β-galactosidase (SA-β-gal) activity and the exit of HMGB1 from the nucleus in Setd8-LKO hepatocytes (Fig. 4B). Interestingly, HMGB1 was also detected in endonucleotic vesicles (Fig. 4B). In addition, H3K9Me_3_ and DAPI staining revealed large focal accumulations, reminiscent of senescence-associated heterochromatin foci (SAHFs) (fig. S3D). Importantly, transcriptome analysis of Setd8-LKO cells revealed distinct gene signatures corresponding to hallmark G2/M checkpoint genes, cellular senescence-related and senescence-associated secretory phenotype (SASP) genes (fig. S4A and B and D and E). Significant gene signatures included the group of oncofetal genes (fig. S4C). These genes are highly expressed in embryonic livers, completely silenced after birth and reactivated in hepatocellular carcinoma, conveying “stemness” properties to cancer cells (*24–26*). This latter feature is similar to the recently reported chemotherapy-induced Senescence-Associated-Stemness (SAS) phenotype in lymphomas, which is considered as an early condition triggering escape from cell-cycle arrest and driving more aggressive tumor growth (*27–29*). Taken together, these results suggest that TCPOBOP-treated Setd8-deficient cells exit the G0-phase and replicate DNA. They become growth-arrested at the next G2-phase of the cell cycle but continue DNA replication, without cell duplication, which results in hyperploidy. These cells enter a specialized senescent stage, displaying senescence-specific transcriptome profiles and SAS features.

### Partially preserved chromatin domains in hyperploid cells

In mammals, distinct regions of the genome are tightly associated with the nuclear lamina. These large Lamin-Associated Domains (LADs) play important role in heterochromatin-mediated gene silencing and the overall 3D organization of the genome (*30–31*). Therefore, increased ploidy and increased nuclear membrane surface at the nuclear periphery and the endonucleotic vesicles may affect LAD distribution and chromatin domains. CUT&Tag assays using a lamin A/C antibody revealed minor changes in LADs between wild-type and TCPOBOP-treated Setd8-LKO cells (Fig. 4C and E), suggesting that the same specific genomic regions interact with nuclear envelope-associated or vesicle-associated lamins even in the hyperploid state.

Interestingly, we found a significant overlap between H4K20Me_3_-decorated heterochromatin and LADs (Fig. 4C and D). H4K20Me_3_-decorated heterochromatin in TCPOBOP-treated Setd8-LKO cells remained stable in most regions but a significant number of new sites were also detected (Fig. 4F). The association of H4K20Me_3_ with nuclear lamins at the inner nuclear envelope and the outer membrane of the internalized vesicles was further validated by proximity ligation assays (PLA) (fig. S5).

Most of the open chromatin areas detected in the wild-type genome, including H3K27-acetylated nucleosomes and Transposase-accessible regions were unaffected in TCPOBOP-treated Setd8-LKO cells (fig. S6). Despite the number of new locations identified in Setd8-LKO cells, the overall patterns and the periodic clustering of the H3K27ac-decorated, transposase-accessible euchromatin domains and those of H4K20Me3-occupied, lamina-associated heterochromatin domains were found to be largely preserved (fig. S6).

Examination of the binding locations of the architectural transcription factor CTCF (*32*) and the cohesion ring complex component Smc3 (*33*) revealed a number of new sites in the hyperploid Setd8-LKO hepatocytes compared to wild-type cells. The new CTCF and Smc3 locations outnumbered those detected in wild-type cells by 3-fold and 1.2-fold, respectively (fig. S7A and 8A and D). However, the majority of the new CTCF locations did not possess a canonical CTCF binding sequence motif and had lower binding strength and a more diffuse distribution (fig. S7B and C). Smc3 binding in the new locations had a lower affinity and a greatly altered distribution relative to CTCF-occupied putative loop-anchor positions (fig. S8B and C). The results suggest that in Setd8-LKO hepatocytes, CTCF and Smc3 associate with genomic sites observed in wild-type cells (constant peaks) but also bind non-specifically into a large number of other regions, probably due to the presence of an abnormally high number of chromosomes. Given the low levels of co-occupancy at canonical CTCF locations, we estimate that the new associations are probably unstable and non-functional.

### Reversal of endonucleosis and hyperploidy

Despite the very dramatic phenotype of TCPOBOP-treated Setd8-LKO mice, with 90% of hepatocytes lost and the remaining cells becoming senescent, the mice were viable for a long period of time after the single TCPOBOP treatment. In fact, similarly to the untreated Setd8-LKO mice (*11*), they developed spontaneous liver cancer at 6 to 8 months of age. We therefore examined the cells beyond the 24-hour time point, for a period of 65 days after the single TCPOBOP injection. On days 7 and 10 we observed a significant reduction in the number of cells containing large vesicles and the parallel appearance of smaller vesicles (Fig. 5A). The smaller vesicles became predominant in most cells on 10 or 20 days after treatment, and accumulated along the nuclear envelope. Importantly, the smaller vesicles did not contain albumin or α-tubulin at these time points, suggesting that cytoplasmic proteins were degraded (fig. S9E). Twenty and 30 days after treatment, numerous cells had nuclear buddings wrapped internally by lamina and micronuclei-like detachments of DNA-containing structures in the cytoplasm (Fig. 5A and G and H). Thirty to 65-days after treatment, intranuclear vesicles were essentially absent and the most common features were segregating nuclei, without mitotic structures, which could be detected by the patterns of DAPI and α-tubulin staining (Fig 5A and fig. S9E). Close-up images captured fission of large intra-nuclear vesicles into microvesicles (Fig. 5B), line-up of microvesicles along the nuclear envelope (Fig. 5C and D), fusion of microvesicles with nuclear membrane exposing the vesicle interior towards the cytoplasm (Fig. 5E and F), budding and detachment of micronuclei (Fig. 5G and H) and various stages of nuclei segregating into smaller-sized ones (Fig. 5I and J). In parallel, we observed a significant reduction in the average nuclear size and an increase in the overall number of cells per field (fig. S9B and D). The PI staining patterns of the nuclei demonstrated a significant reduction in ploidy, with signs for the appearance of cells with aneuploidy (fig. S9a). Ki67-positive cells were greatly reduced and essentially disappeared by day 30 after TCPOBOP treatment (fig. S9C).

**Fig. 5.**
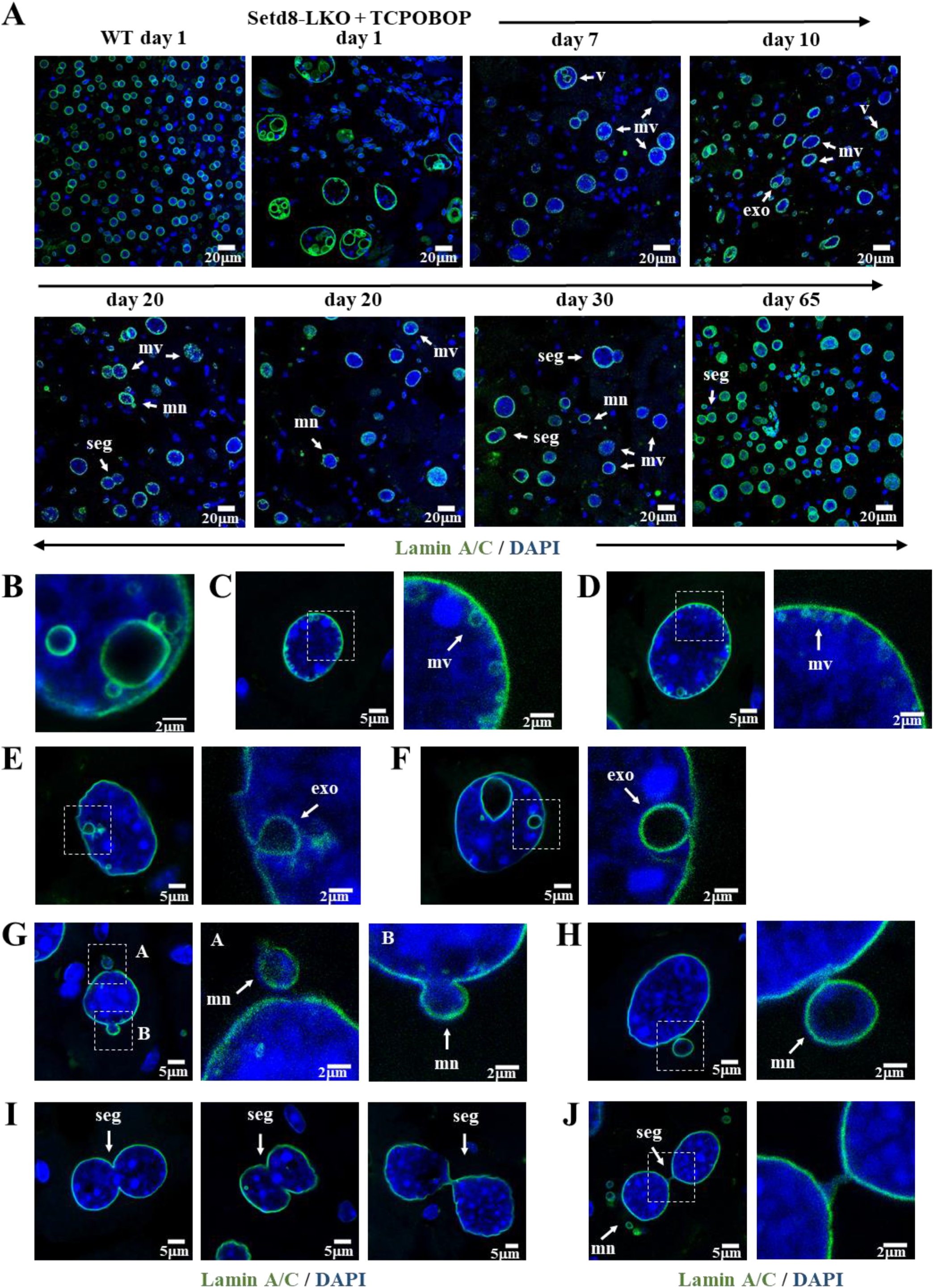
Progressive loss of endonucleotic vesicles and segregation-mediated reduction of nuclear size at later periods. (**A**) Representative images of Lamin A/C-stained hepatocyte nuclei in untreated wild type (WT day 1) and TCPOBOP-treated Setd8-LKO mice 1, 7, 10, 20, 30 and 65 days after the treatment. Arrows indicate: v, large vesicles; mv, microvesicles; exo, exonucleotic vesicles; mn, micronucleus-like structures; seg, segregating nuclei. (**B**) Close-up image of separating microvesicles from a larger endonuleotic vesicle. (**C-D**) Microvesicles accumulating and fusing to the nuclear lamina. (**E-F**) Exonucleotic-like structures releasing vesicle content to the cytoplasm. (**G-H**) Nuclear budding and micronucleus-like structures. (**I**) Segregating nuclei. (**J**) Segregating and budding nuclei.

Taken together, the above results suggest that the processes leading to increased nuclear size, hyperploidy, and accumulation of endonucleotic vesicles in senescent cells are reversible.

## Discussion

Setd8 deficiency leads to genome instability, defects in mitotic chromatin compaction, cell cycle arrest (1–9) and metabolic reprogramming (*12*), which eventually culminate in necrotic cell death (*11*). These effects are in apparent contradiction to the long-term survival phenotype and the late-onset development of liver cancer in liver-specific Setd8 knock-out mice (*11*). To reconcile this paradox, we analyzed the early stages of cell division-dependent hepatocyte death in Setd8-LKO mice.

We found that under mitogenic stress conditions, while most hepatocytes die via necrosis within 24 hours, approximately 10% of the cells escape necrotic death by entering a senescent state. Although, surviving Setd8-LKO hepatocytes displayed most of the bona fide characteristics of senescent cells (*22–23*), including cell cycle withdrawal, DNA damage, senescence-related transcriptome changes, SA-β-galactosidase activity, SASP production and nuclear enlargement, they also exhibited transient hyperploidy and endonucleosis, not normally seen in senescent cells. Our results provide mechanistic explanations for the development of the above unusual phenotype and support a model, schematically presented in fig S10. The first step involves mitogenic stimulus-dependent endoreduplication in the cell division-incompetent, G2-phase arrested Setd8-LKO hepatocytes. Endoreduplication generates cells with multiple copies of each chromosome, which could confer a selective survival advantage by compensating for the detrimental effects of extensive DNA damage, such as transcriptional deregulation, the emergence of lethal loss-of-function mutations and/or haploinsufficency.

Nonetheless the cells, which escape necrosis, enter a senescent state and must cope with the increased nuclear DNA content either by increasing nuclear size or by initiating other structural changes in the nucleus. A plethora of previous studies have established the view that the shape of the nuclear envelope is subject to alterations regulated by internal forces connecting the lamin fibers to chromatin via LADs and external anchoring to microtubules and cytoskeletal filaments by nesprins and SUN domain proteins (*34, 35*). Setd8-deficient hepatocytes have greatly enlarged nuclei and consequently elongated nuclear envelope, with increased elasticity. We observed thinner internal lamin fibers in Setd8-LKO cells and a proportionally increased lamin protein content per unit areas, which is indicative of a mechanical stretching of existing and new lamin polymers. The resulting elasticity of the nuclear envelope may not only allow for an increase in nuclear size, but may also counterbalance potential mechanical force-driven adverse effects caused by hyperploidy. We also found that LADs are enriched with H4K20 trimethylated nucleosomes. Although Setd8-deficient cells cannot mono-methylate H4K20, the existing H4K20Me_1_ histone tails can be up-methylated to di-and trimethylated forms by Suv4-2h1/2 (*4, 5*). Consistent with this, we observed elevated levels of H4K20Me_3_ in Setd8-LKO mice, which can efficiently associate with the nuclear lamina and generate an additional internal force that folds the elastic nuclear envelope inwards.

Folding of the lamina towards the nuclear interior and the subsequent membrane closure, generates intranuclear vesicles with entrapped cytoplasmic proteins, a process we term endonucleosis. Our results also demonstrate that the parallel activation and function of the cytoplasmic autophagy machinery, which is induced by elevated AMPK levels in Setd8-LKO mice (*12*), is a prerequisite for facilitating and completing the endonucleosis process. Given the context and the conditions under this process takes place, it is tempting to speculate that endonucleosis represents an additional survival mechanism to counteract the extreme size expansion of the cells. Internalization of cytoplasm via autophagy could contribute to the elimination of toxic substances and structures that accumulate in the cytoplasm and to the preservation of the overall nucleus/cell size ratios within a range that is compatible with survival.

In line with this scenario, we found that the unusual senescent phenotype characterized by hyperploidy and endonucleosis corresponds to an unstable and transient cellular state. A few days after their appearance, the endonucleotic vesicles became smaller and gradually disappeared. In parallel, the nuclear size and cellular ploidy were also reduced via mechanisms involving the segregation of the existing nuclei without mitosis and the extensive budding and detachment of smaller nuclear structures, reminiscent of micronuclei (*36, 37*). While these processes may act towards normalizing the physical characteristics of the surviving cells and their genome, they may also generate aneuploidy, which together with the activation of oncofetal genes and the parallel accumulation of inflammatory cells, could provide a fertile ground for the initiation of hepatocellular carcinoma at later stages.

Taken together, the senescence features described here, provide support for the concept of non-genetic function of the genome and chromatin (*38*) by demonstrating their importance in the regulation of nuclear envelope plasticity in cellular survival under stress conditions.

## Materials and Methods

### Mice

Setd8^lox/lox^-AlbCre (Setd8-LKO) mice were generated by crossing mice carrying floxed exon 7 allele with Alb-Cre transgenic mice (11). Hepatocyte-specific, full inactivation of Setd8 in these mice was observed from postnatal day 20 and onwards (11). LC3-GFP transgenic mice (19) (RBRC00806) and mice carrying floxed alleles of Atg5 gene (21) (B6.129S-Atg5 [tm1Myok]) were from RIKEN BioResource Center and were obtained from Georgios Chamilos (Univ. of Crete). Setd8/Atg5-LKO double knockout mice were obtained by crossing Setd8^lox/lox^-AlbCre mice with Atg5^lox/lox^ mice. The resulting single and double knockout mice were fertile and viable over 12 months of age. Mice were kept in grouped cages in a temperature-controlled, pathogen-free facility on a 12 hr light/dark cycle and fed with standard chow diet containing 19% protein and 5% fat (Altromin 1324) and water ad libitum. All animal experiments were approved by the Ethical Review Board of IMBB-FORTH and the Animal Ethics Committee of the Prefecture of Crete and were performed in accordance with the respective national and European Union regulations. All experiments were performed in randomly chosen age-matched male mice. No blinding was used in this study. Unless otherwise indicated, mice were treated and analyzed at 45-50 days after birth. Treatment of mice were performed with a single intraperitoneal injection of either vehicle (corn-oil) or 3 mg/kg 1,4-bis-[2-(3,5,-dichloropyridyloxy)] benzene (TCPOBOP, T1443, Sigma-Aldrich).

### Hematoxylin and Eosin (H&E) staining and immunofluorescence assays

Fixation, paraffin infiltration and H&E staining of liver tissues were performed as described previously (25). For immunofluorescence staining freshly isolated liver tissues were either prefixed in 4% paraformaldehyde (PFA) in PBS for 2 hours and treated sequentially with 15% and 30% sucrose, before embedding in Optimal Cutting Temperature (OCT) embedding medium or embedded unfixed and were frozen in liquid nitrogen. Cryosections of pre-fixed liver tissue (5 to 7 μm thick) were boiled in 10 mM tri-sodium citrate buffer pH 6 with 0.05% Tween-20 twice for 5 minutes for heat-induced epitope retrieval. Frozen sections of unfixed tissue (5 to 7 μm thick) were air dried and fixed in 4% formaldehyde for 15 min at room temperature, blocked in 5% BSA in PBS containing 0.3% Triton X-100 for 1 hour and then incubated with 1% BSA, 0.3% Triton X-100 in PBS with the indicated primary antibodies at 4^0^C overnight. After incubation with AlexaFluor 594- or AlexaFluor 488-conjugated goat anti-rabbit or anti-mouse secondary antibodies for 1 hour at room temperature and counterstained with 1 μg/ml DAPI for 10 min, the slides were covered with Mowiol® 4-88 Reagent (EMD Millipore, 475904). Fluorescence images were observed using a Leica TCS SP8 confocal microscope. Image analysis was performed with the FIJI/Image J software.

For chromosomal ploidy evaluation, isolated nuclei were seeded onto Superfrost® Plus glass microscope slides, fixed with 70% EtOH and stained with 50μg/ml propidium iodide (Sigma-Aldrich, P4864) for 30 minutes. After washing with PBS and mounting with Mowiol® 4-88 Reagent, PI images were collected using Operetta (Perkin Elmer) high content microscope and PI intensity data were evaluated using the Harmony Software 4.1 of PerkinElmer.

### High Content Microscopy analyses of wide field and confocal images

Tissue cryosections and isolated nuclei on slides labeled with immunofluorescence were captured with a 20x or 40x lens (Olympus–Shinjuku City, Tokyo, Japan) with the Operetta High Content Screening Microscope (HCSM) (PerkinElmer, Waltham, MA, USA) in wide field and analyzed using Harmony software 4.1 with PhenoLOGIC (PerkinElmer) or the open source software FIJI/Image J.

Segmentation of the nuclei was based on DAPI nuclear dye. Nuclei that were not intact, i.e. with no clear boundaries, or nuclei that were interrupted by the field boundaries were excluded from the analysis. Only whole, well-formed round nuclei were used for the analysis while non-hepatocyte cells were excluded based on their elongated shape and smaller size. For the measurement of nuclear size, the selected segmented nucleus area was measured in μm^2^ in thirty different tissue sections and 5000 randomly selected nuclei were plotted in GraphPad Prism 6. For the determination of the relative protein expression levels, the sum intensity of selected nuclei was calculated and plotted in GraphPad Prism 6. In the cases where background fluorescent noise was observed, this was estimated per nuclear area and subtracted from the nuclear sum intensity prior to plotting the graph. For the estimation of the number of nuclei per field in different time points, only fields with tissue coverage of 85 to 86% were selected.

For the construction of ploidy graphs, nuclei were captured with the 20x lens in the HCSM. Nuclei segmentation was based on the Propidium Iodide (PI) dye that quantifies the nuclear DNA content. Aggregates were excluded from the analysis. The PI sum intensity of each nucleus was calculated and was plotted in Microsoft Excel where the Y axis in the graphs represents the cell count within a range of sum intensities using a standard increase of 99999.

For the nuclear lamina intensity estimation, whole tissue cryosections were captured with a Leica TCS SP8 confocal microscope with a 63x lens and analyzed with using Fiji/Image J. One plane from the stack was selected, and the mean intensity of an area, delimited by tracing a two-pixel width line along the lamin-stained membrane over the point with the highest fluorescent intensity, was estimated. Measurements on a given nucleus were repeated multiple times and in multiple areas to minimize technical errors and the values were averaged per nuclei and plotted in Graphpad Prism 6. For estimation of nuclear lamina thickness, the function Plot Profile in FIJI/Image J was used, and the images were analyzed with the parameter setting ‘full width at half-maximum’, that gives the distance between the two intensity points in a selected area. For each nucleus, multiple measurements were performed in the same area. Areas that were not clear or the nuclear membrane was folded, were excluded from the measurement.

For the estimation of nuclear/cell area ratios, images from E-cadherin stained liver sections were evaluated by using FIJI/ImageJ analysis software. The perimeter of each cell and nucleus were marked by the Polygon selection tool. After clearing the background, nuclear and cellular areas were measured by the Analyze Particles tool using minimum area threshold of 50 μm^2^ for wild type and 300 μm^2^ for Setd8-LKO.

### Preparation of nuclei

Hepatocyte nuclei were prepared as follows: liver tissue was minced into small pieces in 20 volumes of ice-cold buffer containing 10 mM Tris pH 7.5, 2 mM MgCl_2_, 3 mM CaCl_2_ and protease inhibitor cocktail (Complete^TM^ EDTA-free, Roche), and homogenized by 10 strokes using Dounce device. The suspension was filtered through 100 μm mesh strainer and centrifuged for 5 min at 400g, at 4^0^C. The resulting pellet was resuspended in a buffer containing 10 mM Tris pH 7.5, 2 mM MgCl_2_, 3 mM CaCl_2_, 10% glycerol, 1% NP40 and protease inhibitor cocktail (Complete^TM^ EDTA-free, Roche). After incubation in ice for 3 min the nuclei were collected by centrifugation for 5 min at 400g, at 4^0^C, washed twice with PBS and used for downstream applications.

### Electron microscopy

Liver tissues were fixed for 2 hours at room temperature in 0.08 M sodium cacodylate buffer, containing 2% glutaraldehyde and 2% paraformaldehyde. After a 1-hour post-fixation step with 1% osmium tetroxide, the slides were treated with 1% uranyl acetate for 20 min. The samples were dehydrated with serial ethanol treatment and subsequently embedded in Durcupan resin/propylene oxide (Polysciences). Approximately 100 nm thin sections on copper grids were observed at 80 kV with a JEOL JM2100 transmission electron microscope.

### DNA Fluorescence *in situ* hybridization (DNA-FISH)

BAC clones were obtained from BACPAC Resource Center CHORI. The following BAC clones were used for probe preparation RP24-353H5 spanning *Afp* locus in chromosome 5; RP24-82E10 spanning *Pkm* locus in chromosome 9; RP24-389I13 for *Igf1* locus in chromosome 10.

BAC clones were labeled by nick-translation using Aminoallyl-dUTP-Cy3 (Jena Bioscience). 100 ng of the labeled probes were mixed with 1μg COT-1 DNA (Invitrogen), lyophilized and resuspended in formamide. After denaturation at 95^0^C for 10 min, the probes were diluted to prepare hybridization buffer containing 2xSSC, 10% Dextran Sulphate and 50 mM Na_2_HPO_4_.

18 μm thin liver cryosections were treated with 10 mM Na_3_-citrate pH 6.0 for 10 min at 80^0^C and placed to room temperature for 1 hour. The slides were sequentially treated with 70%, 80%, 95% and 100% ethanol for 3 min each. After DNA probe addition, the slides were incubated at 80^0^C for 5 min for additional denaturation, followed by hybridization at 37^0^C for 14 hours. The slides were washed first by 0.5x SSC/0.1% Tween-20 for 5 min at 70^0^C, followed by washing with 1x SSC/0.1% Tween-20 for 5 min at room temperature, and by 4x SSC/0.1% Tween-20 for 5 min, by 2x SSC for 5 min and finally by PBS. The slides were further treated with 0.5% Triton X-100 in PBS for 24 hours at 4^0^C, counterstained by DAPI and mounted using Prolong Gold Antifade reagent (Invitrogen).

### Senescence-associated beta-galactosidase (SA-β-gal) and PAS staining

Liver sections from OCT-embedded frozen livers were fixed in 0.2% glutaraldehyde for 10 min. The sections were washed with PBS and stained for 2 hours with a solution containing 0.5 mg/ml X-Gal, 5 mM of K3Fe(CN)6, 5 mM K4Fe(CN)6, 2 mM MgCl_2_, 150 mM NaCl, 40 mM citric acid/sodium dihydrogen phosphate, pH 4. The sections were counterstained with eosin for 1.5 min and observed with light microscopy using Olympus CX23T microscope. Images were taken using Axiocam ERc5s (ZEISS) and the ZEN microscopy software (ZEISS).

Periodic-acid Schiff (PAS) staining was performed in formalin fixed paraffin-embedded liver sections. The sections were treated with 0.5% Periodic acid for 5 min and stained in Schiff’s reagent (Merck) for 10 min. After washings with PBS, the sections were counterstained with hematoxylin and observed by light microscopy as above.

### Proximity Ligation Assay (PLA)

Proximity Ligation Assays were performed in liver cryosections from frozen tissue using Duolink® In situ Red Starter Kit Mouse/Rabbit from Sigma-Aldrich, according to the manufacturer’s instructions. The pairs of primary antibodies were mouse anti-LaminA/C (Cell Signaling Technology, #4777), with rabbit anti-LaminB1 (Abcam, ab16048); and mouse anti-LaminA/C (Cell Signaling Technology, #4777) with rabbit anti-H4K20Me_3_ (Abcam, ab9053). Following PLA reaction, the slides were counterstained with 1 μg/ml DAPI for 10 min and observed in Leica TCS SP8 confocal microscope.

### CUT&Tag and ATAC-seq assays

CUT&Tag assays were performed as follows: 10^5^ nuclei were resuspended in a buffer containing 20 mM HEPES–KOH pH 7.9, 10 mM KCl, 0.1% Triton X-100, 20% Glycerol, 0.5 mM Spermidine, protease inhibitor cocktail (Complete^TM^ EDTA-free, Roche) and incubated for 10 min with Concanavalin A-conjugated magnetic beads (Epicypher) that were previously activated by repeated washings with a buffer containing 20 mM HEPES pH 7.9, 10 mM KCl, 1 mM CaCl_2_, 1 mM MnCl_2_. The reactions were supplemented with 0.5 volume of a buffer containing 20 mM HEPES, pH 7.5, 150 mM NaCl, 0.5 mM Spermidine, protease inhibitor cocktail, 0.01% Digitonin, 2 mM EDTA and 0.5 μg of the anti-Lamin A/C or anti-H4K20Me_3_ or IgG negative control primary antibodies. The samples were incubated overnight at 4^0^C by gentle agitation. The beads were collected and supplemented with a buffer containing 20 mM HEPES, pH 7.5, 150 mM NaCl, 0.5 mM Spermidine, protease inhibitor cocktail, 0.01% Digitonin and 0.5 μg of anti-rabbit or anti-mouse secondary antibodies (Epicypher). After incubation for 30 min at room temperature, the beads were washed twice with the same buffer and resuspended in the same buffer containing 300 mM NaCl. Protein A/G-fused Tn5 transposase protein (CUTANA pA/G-Tn5, Epicypher) was added and the samples were incubated for 1 hour at room temperature followed by washing with a buffer containing 20 mM HEPES, pH 7.5, 300 mM NaCl, 0.5 mM Spermidine, protease inhibitor cocktail and 0.01% Digitonin. The beads were collected and resuspended in the above buffer containing 10 mM MgCl_2_, and incubated for 1 hour at room temperature. The beads were collected and washed once with 10 mM TAPS pH 8.5 and 0.2 mM EDTA. DNA was released from the beads by incubation in 10 mM TAPS pH 8.5 and 0.1% SDS for 1 hour at 58^0^C. After quenching by the addition of 3 volumes of 0.67% Triton-X100, the samples were used for PCR reaction (15 cycles) using universal i5 primers and barcoded i7 primers. After cleanup with AMPure beads (Beckman Coulter), quantification and quality evaluation in Agilent Bionalyzer, the libraries were sequenced using Illumina NextSeq500 platform.

For ATAC-seq reactions we used the ATAC-seq kit from Active Motif. Tagmentation reactions using 10^5^ nuclei, DNA purification and PCR amplifications using combinations of indexed i5 and i7 primers were performed according to the manufacturer’s instructions. The resulting libraries were sequenced using Illumina NextSeq500 platform.

### Chromatin Immunoprecipitation (ChIP) assays

ChIP assays were performed as described previously (26, 41). Briefly, livers were minced to small pieces and crosslinked with 1% formaldehyde for 10 min. After the addition of glycine at 0.125 M final concentration, the cross-linked cells were washed with a buffer containing 50 mM HEPES pH 7.9, 100 mM NaCl, 1 mM EDTA, 0.5 mM EGTA and sequentially treated with a buffer containing 0.25% Triton-X100, 10 mM EDTA, 0.5 mM EGTA, 20 mM HEPES pH 7.9 and a buffer containing 0.15M NaCl, 1 mM EDTA, 0.5 mM EGTA, 20 mM HEPES pH 7.9 for 10 minutes each. The resulting crosslinked nuclei were resuspended in 10 volumes of 50 mM HEPES pH 7.9, 140 mM NaCl, 1 mM EDTA, 0.1% Na-deoxycholate, 0.5% Sarkosyl, and protease inhibitor cocktail (Complete^TM^ Roche), sonicated for 10 minutes in Covaris Sonicator instrument with maximum setting. After sonication, the samples were supplemented with 0.5 volume of 50 mM HEPES pH 7.9, 140 mM NaCl, 1 mM EDTA, 3% Triton X-100, 0.1% Sodium deoxycholate, and protease inhibitor cocktail, and centrifuged for 15 minutes at 20.000g. The soluble chromatin was incubated overnight with Dynabeads Protein G (Invitrogen), that were prebound by 5 μg of the primary antibodies recognizing H3K27ac-modified histones, CTCF and Smc3. The beads were washed twice with a buffer containing 50 mM HEPES pH 7.9, 140 mM NaCl, 1 mM EDTA, 1% Triton X-100, 0.1% Na-deoxycholate, 0.1% SDS, 0.5 mM PMSF, 2 μg/ml aprotinin, once with the same buffer but containing 500 mM NaCl, once with a buffer containing 20 mM Tris, pH 8.0, 1 mM EDTA, 250 mM LiCl, 0.5 % NP-40, 0.5 % Na-deoxycholate, 0.5 mM PMSF, 2 μg/ml aprotinin and once with a buffer containing 10 mM Tris pH 8.0, 0.1 mM EDTA, 0.5 mM PMSF, 2 μg/ml aprotinin. Immunoprecipitated chromatin was eluted from the beads and reverse crosslinked by overnight incubation in a buffer containing 50 mM Tris-HCl, pH 8.0, 1 mM EDTA, 1% SDS and 50 mM NaHCO_3_ at 65^0^C. The samples were then incubated with 25 μg/ml RNase-A for 30 minutes at 37^0^C, followed by incubation with 50 μg/ml Proteinase-K for 2 hours at 42^0^C. DNA was extracted by phenol/chloroform and precipitated with ethanol.

Libraries were prepared from 10 ng input DNA NEBNext Ultra II DNA Library Prep *Kit* of New England Biolabs according to the manufacturer’s instructions and sequenced in Illumina NextSeq500 sequencer.

### CUT&Tag, ATAC-seq and ChIP-seq data analysis

Computational analyses of genome-wide mapping data were performed as described previously (25, 41), with minor modifications. Briefly, before performing any alignment or additional analysis, the quality of all the FASTQ files and the reads they contain was assessed using *FastQC* https://www.bioinformatics.babraham.ac.uk/projects/fastqc. To remove low-quality bases and adapter sequences from the raw FASTQ files, we employed *Trimmomatic* (version 0.39) (42). Duplicate reads were identified and removed using *samtools* version 1.10 (43). The FASTQ files were mapped on the UCSC mm10 genome using *hisat2* with default settings (44). We retained only the reads with a mapping quality score greater than 30 and removed blacklisted regions from our BAM files which are known to be susceptible to technical artifacts and spurious signals. The resulting BAM files were converted to bigwig files using *deeptools* (version 3.3.2) (45) and visualized in the UCSC Genome Browser.

Before identifying peak regions and to account for variations in the number of sequencing reads between ChIP and CUT&Tag samples, we standardized the total read count for each sample by equally reducing reads proportionally based on the sample with the fewer reads. Peaks were called using *MACS2* (2.2.7.1) for mouse genome with a *pvalue cutoff* of 1.00e-13(2.2.7.1) (46). In the case of Lamin A/C and H4K20Me_3_ CUT&Tag data, peak calling was performed by SICER2 (47). De novo motif search was run using HOMER45 within the median +/- nt intervals of each sample separately around the peak summit of 25% best scoring (pvalue) peaks.

To gain insightful visual representations of the genomic binding patterns and chromatin profiles, we employed *deeptools* (version 3.3.2) utilizing the “computeMatrix” module and calculated the read density at the peak centered regions. Venn diagrams of overlapping regions between conditions were created using custom made scripts in R version (4.3.0).

### RNA purification and RNA-sequencing

Total RNAs were purified from liver tissues by brief homogenization in 10 volumes of Trizol reagent using Polytron device, followed by incubation at room temperature for 5 min and the addition of 0.2 volumes of chloroform and further incubation at room temperature for 3 minutes. The samples were centrifuged at 12000 g for 15 minutes at 4^0^C and the aqueous phase was collected and was precipitated by the addition of equal volume of isopropanol. After 10 minutes at room temperature, the RNA was collected by centrifugation at 12000g for 15 minutes. The pellet was resuspended in H_2_O and re-precipitated with ethanol. The RNA samples were further purified by digestion with 10 units of DNase-I for 10 min at 37^0^C, followed by purification with phenol/chloroform extraction and ethanol precipitation.

RNA-seq libraries were generated using NEBNext Ultra II RNA Library preparation kit from New England Biolabs and sequenced in an Illumina NextSeq 500 system. Raw sequence data were quality assessed and pre-processed using PRINSEQ version 0.20.4 (48). Pre-processed FASTQ files were subsequently mapped to the UCSC mm10 reference genome, using HISAT2 version 2.1.0 with the argument –score-min L,0.0–0.5, setting the function for the minimum valid alignment score. The resulting BAM files were analyzed with the Bioconductor package metaseqR (49). Differential gene expression analysis and visualizations were performed as described previously (25). Gene Set Enrichment Analysis (GSEA) were performed using the GSEA software version 3.0 (50), with default parameters and FDR<0.25 cutoff. The analysis was performed on the RPGM normalized expression values of the differentially expressed genes.

### Quantification and statistical analysis

Comparisons between two groups were performed using Student unpaired t test. All statistical analyses were performed with Graphpad Prism 6 to 8 (GraphPad).

Sample size was determined empirically based on similar studies with Setd8-LKO mice (Refs. 11 and 12). No statistical method was used to predetermine sample size. As a rule, all of the imaging-based assays were performed at least 2 different dates with 3 different biological replicates (mice) each time and at least 8-10 technical replicates (sections) each time. RNA-seq analyses involved RNAs from 4 and 5 biological replicates collected at the same date. ChIP-seq and CUT&Tag assays involved at least 2 biological replicates.

### Antibodies

Antibodies were used for Immunofluorescence staining (IF) in the indicated dilutions, for CUT&Tag or for Chromatin Immunoprecipitation (ChIP) from the following vendors:

#### Cell Signaling technologies

Mouse anti-Lamin A/C (4C11), #4777, RRID:AB_10545756 (IF: 1:100, CUT&Tag)

Mouse anti-α-Tubulin (DM1A), #3873, RRID:AB_1904178 (IF: 1:400)

Rabbit anti-FANCD2 (D5L5X), #16323, RRID:AB_2798761 (IF: 1:100)

Rat anti-RPA32/RPA2 (4E4), #2208, RRID:AB_2238543 (IF: 1:50)

Rabbit anti-Phospho-Histone H2A.X (Ser139) (20E3), #9718, RRID:AB_2118009 (IF: 1:200)

Goat anti-mouse IgG (H+L), F(ab’)2 Fragment (Alexa Fluor® 594 Conjugate), #8890, RRID:AB_2714182 (IF: 1:500)

Goat anti-rabbit IgG (H+L), F(ab’)2 Fragment (Alexa Fluor® 488 Conjugate), #4412, RRID:AB_1904025 (IF: 1:500)

Goat anti-rabbit IgG (H+L), F(ab’)2 Fragment (Alexa Fluor® 594 Conjugate), #8889, RRID:AB_2716249 (IF: 1:500)

Goat anti-mouse IgG (H+L), F(ab’)2 Fragment (Alexa Fluor® 488 Conjugate), #4408, RRID:AB_10694704 (IF: 1:500)

#### Abcam

Rabbit anti-LaminB1, ab16048, RRID:AB_443298 (IF: 1:100)

Rabbit anti-SUN2 (EPR6557), ab124916, RRID:AB_10972497 (IF: 1:100)

Rabbit anti-SMC3, ab9263, RRID:AB_307122 (IF: 1:800, ChIP)

Rabbit anti-Histone H4 (tri methyl K20), ab9053, RRID:AB_306969 (IF: 1:200, IF and CUT&Tag)

Rabbit anti-Histone H4 (mono methyl K20), ab9051, RRID:AB_306967 (IF: 1:200)

Rabbit anti-Histone H3 (tri methyl K9), ab8898, RRID:AB_306848 (IF: 1:200)

Rabbit anti-Histone H3 (acetyl K27), ab4729, RRID:AB_2118291 (ChIP:1:100)

Rabbit anti-Ki67, ab15580, RRID:AB_443209 (IF: 1:200)

Rabbit anti-HMGB1, ab18256, RRID:AB_444360 (IF: 1:200)

Goat Anti-Guinea pig IgG H&L (Alexa Fluor® 647), ab150187, RRID:AB_2827756 (IF: 1:200)

#### Novus Biologicals

Rabbit anti-Calnexin, NB100-1974, RRID:AB_10001873 (IF: 1:50)

#### BD Biosciences

Mouse anti-E-cadherin (clone 36), 610181, RRID:AB_397580 (IF: 1:200)

#### Santa Cruz Biotechnologies

Rabbit anti-HNF4α (H-171), sc-8987, RRID:AB_2116913 (IF: 1:100)

#### EMD Millipore

Rabbit anti-CTCF, 07-729, RRID:AB_441965 (ChIP)

#### Progen

Guinea pig anti-p62/SQSTM1 (C-terminus), GP62-C, RRID:AB_2687531 (IF: 1:100)

#### Proteintech

Rabbit anti-Albumin, 16475-1-AP, RRID:AB_2242567 (IF: 1:50)

#### Jackson Immuno Research Laboratory

Alexa Fluor® 594 AffiniPure Donkey Anti-Rat IgG (H+L), 712-585-153, RRID:AB_2340689 (IF: 1:200)

## Acknowledgments

We thank H. Kontaki, M. Koukaki and E. Moltsanidou for technical assistance and discussions during the course of the work; E. Stratidaki, N. Gounalaki and E. Dialynas of the IMBB-Genomics facility for assistance with NGS library preparations and sequencing; S. Papadogiorgaki for help with electron microscopy; M. Vasilarou for initial RNA data analyses; P. Hatzis for editing the manuscript.

## Data and materials availability

All data are available in the main text or the supplementary materials. RNA-seq, ChIP-seq, ATAC-seq and CUT&Tag data are available at the Gene Expression Omnibus (GEO) under accession number GSE242719.

---

**Fig. S1.**
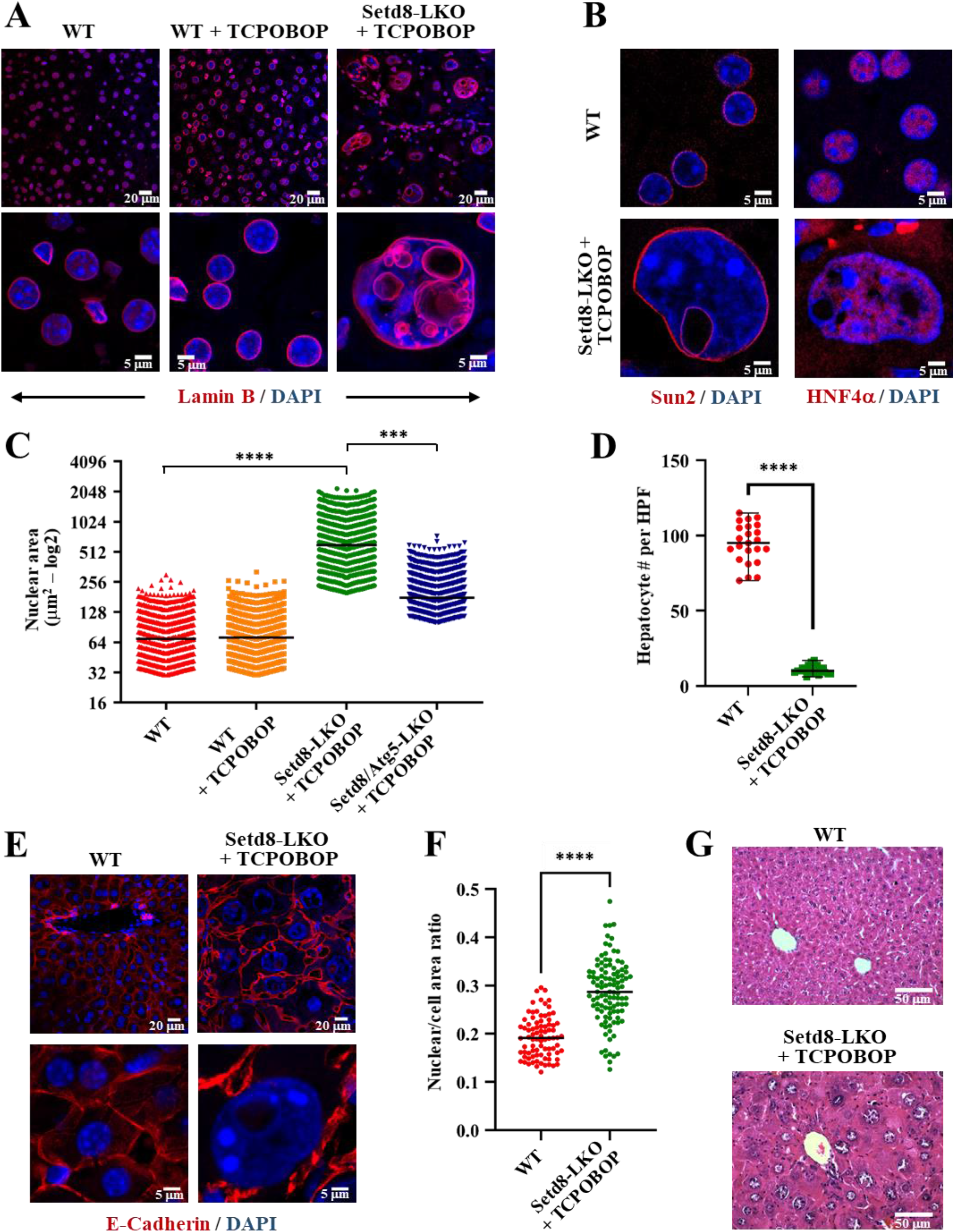
Nuclear envelope composition and characterization of cell and nucleus size in hepatocytes displaying endonucleosis. **(A)** Lamin B antibody immunostaining of liver sections from wild type (WT) mice and wild type or Setd8-LKO mice after 24 hours of treatment with TCPOBOP. Fluorescent images of the cells are shown at two different magnifications. **(B)** Immunofluorescence detection of the inner nuclear envelope component Sun2 and the nuclear protein HNF4α. **(C)** Comparison of the relative nuclear areas. High Content Microscopy measurements were performed in n=4295 nuclei of wild type hepatocytes, n=4476 nuclei of TCPOBOP-treated wild type hepatocytes, n=4095 nuclei of TCPOBOP-treated Setd8-LKO hepatocytes, and n=4091 nuclei of TCPOBOP-treated Setd8/Atg5-LKO hepatocytes. Black horizontal lines indicate median values. **(D)** Quantification of hepatocyte numbers in 20 High Power Fields (HPF) in wild type and TCPOBOP-treated Setd8-LKO livers. Mean values and standard errors are indicated. **(E)** E-Cadherin staining to determine cellular borders and the number of nuclei in hepatocytes. Note the occasional presence of binuclear cells in both wild type and TCPOBOP-treated Setd8-LKO hepatocytes. **(F)** Determination of the relative nuclear size compared to total cell areas in n=110 wild type and n=107 TCPOBOP-treated Setd8-LKO hepatocytes. **(G)** Hematoxylin-eosin (H&E) staining of liver sections from wild type and TCPOBOP-treated Setd8-LKO mice. *** p-value <0.001; ****p-value<0.0001.

**Fig. S2.**
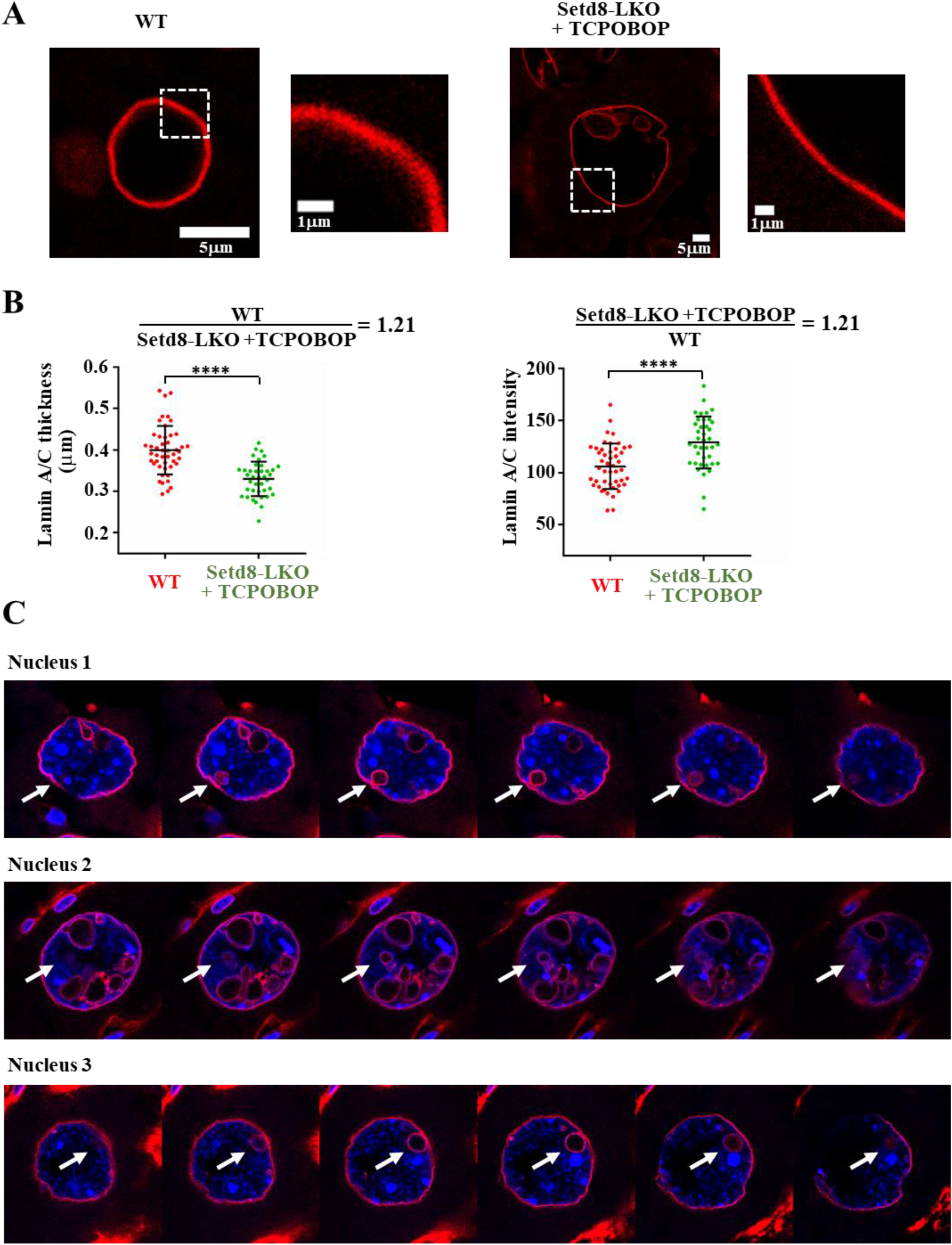
Evaluation of Lamina thickness and the intranuclear positioning of lamin-coated vesicles. **(A)** Representative close-up images of Lamin A/C stained nuclear envelope areas from wild type and TCPOBOP-treated Setd8-LKO hepatocytes. **(B)** Measurements of the thickness and pixel intensity of multiple individual areas of 47 WT and 38 Setd8-LKO hepatocyte nuclei. The same individual areas were used for both thickness and pixel intensity measurement. Note that while the spatial distribution of Lamin A/C signal per unit areas were less dispersed (thinner) in Setd8-LKO nuclei, the average pixel densities were proportionally increased in the same areas. **(C)** Serial confocal z-stack images of nuclei used for 3D reconstruction imaging in Fig. 1B, showing the entirely internal location of multiple vesicles. ****p-value<0.0001.

**Fig. S3.**
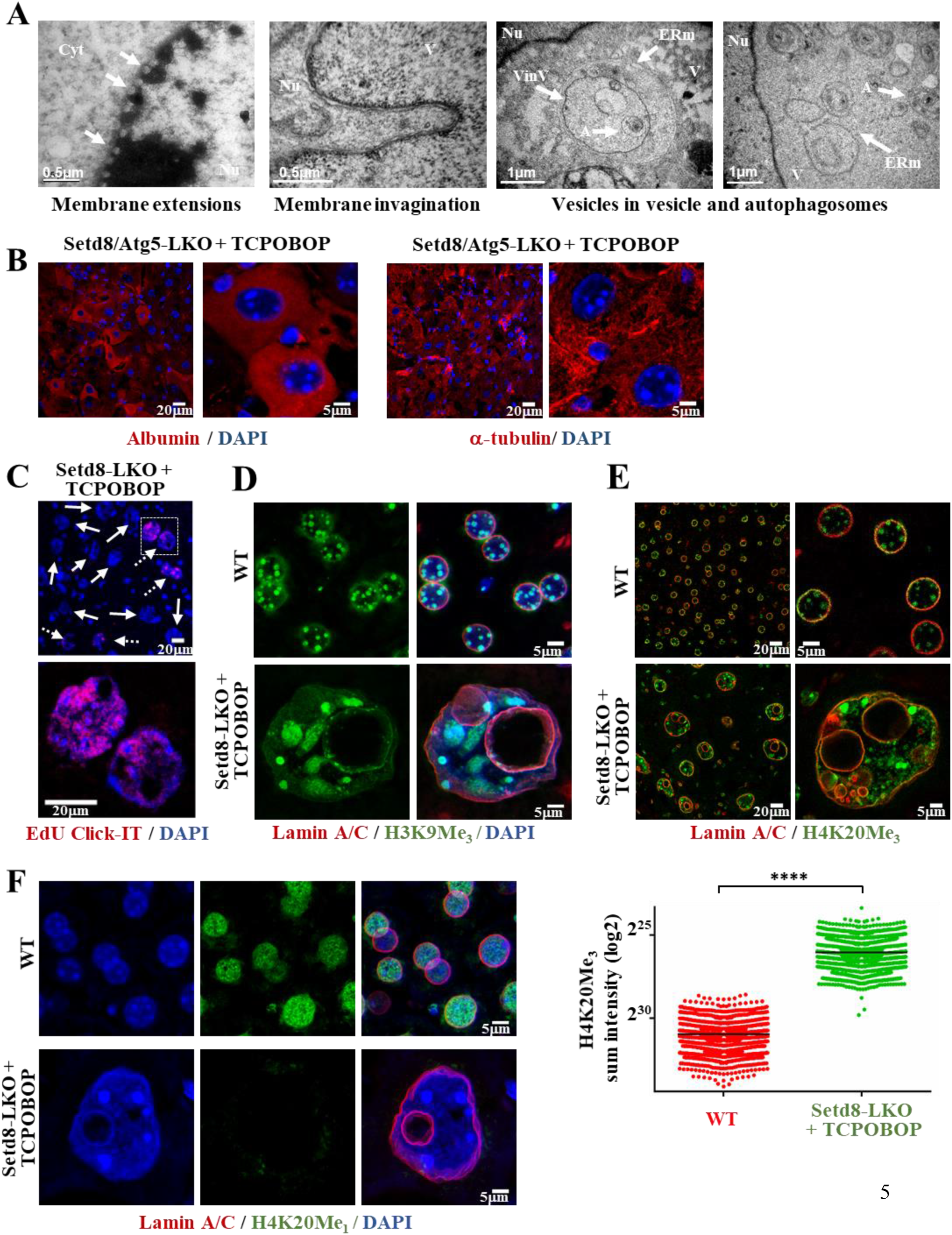
Dynamic reorganization of intranuclear vesicles and histone modification status of cells with endonucleosis. **(A)** Transmission electron microscopic images of TCPOBOP-treated Setd8-LKO hepatocytes capturing disruptions of electron-dense lamina-associated chromatin, indicating membrane extensions (left panel), invagination towards the nuclear interior (2nd panel), vesicles within vesicles (VinV) with autophagosomes (A) and ER membrane (ERm) (3rd and 4th panel). **(B)** Immunostaining of TCPOBOP-treated Atg5/Setd8-LKO, double KO hepatocytes for the Albumin and α-tubulin, demonstrating the lack of endonucleosis-mediated internalization of cytoplasmic proteins in Atg5-deficient cells. **(C)** Fluorescence Click-IT images of liver sections from mice pulse labeled (2 hours) by EdU, showing DNA replication activity in a fraction of TCPOBOP-treated Setd8-LKO hepatocytes at 24 hours after treatment. Panel at right, magnified image of the selected area. Dashed arrows indicate positively stained cells, solid arrows indicate cells that did not incorporate EdU into DNA. **(D)** Immunostaining with antibody recognizing trimethylated H3K9. Note the appearance of large and diffuse DAPI-dense, H3K9Me_3_-labeled heterochromatin foci in Setd8-LKO hepatocytes. **(E-F)** Immunostaining with antibodies recognizing trimethylated **(E)** and monomethylated H4K20 **(F)**. The lower panel of **(E)** shows measurements of H4K20Me_3_ pixel intensities in individual cells. Measurements were from n=2500 wild type and n=2620 TCPOBOP-treated Setd8-LKO hepatocytes. Note the loss of H4K20Me_1_ and the highly increased H4K20Me_3_ levels in Setd8-LKO hepatocytes, indicating up-methylation of existing monomethylated histone H4. ****p-value<0.0001.

**Fig. S4.**
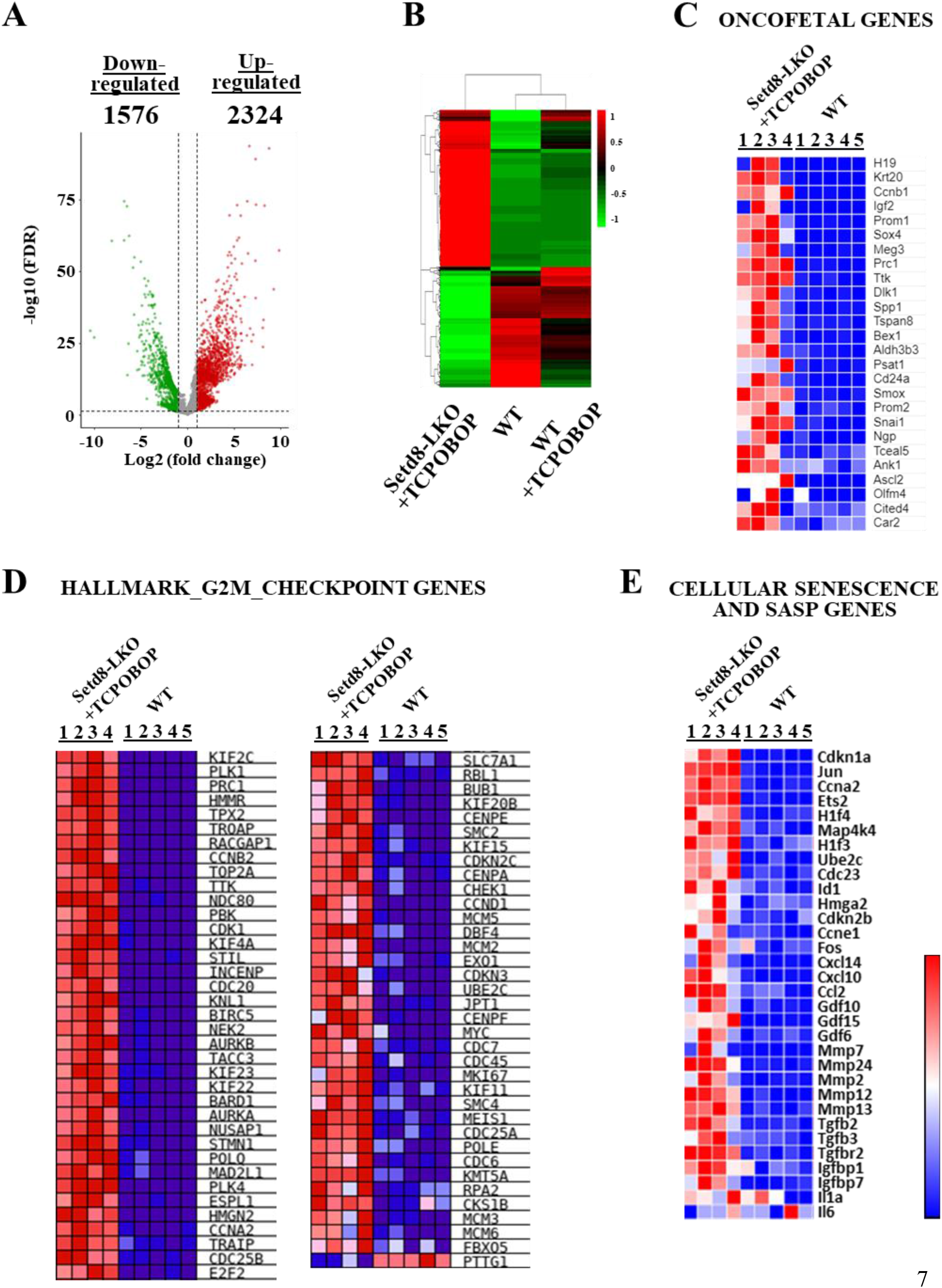
Analysis of global gene expression patterns in wild type and Setd8-LKO hepatocytes. **(A)** Volcano-plot of differentially expressed genes between wild type and TCPOBOP-treated Setd8-LKO hepatocytes. **(B)** Hierarchical clustering analysis and corresponding heatmap of the differentially expressed genes in wild type, TCPOBOP-treated wild type and TCPOBOP-treated Setd8-LKO hepatocytes. Note the distinct profile of the latter, demonstrating Setd8-deficiency-dependent changes. **(C-E)** Heat-map analysis of the normalized RPKM values (raw z-scored) of gene signatures representing hepatic oncofetal genes **(C)**, G2M checkpoint genes **(D)** and senescence-specific genes including SASP genes **(E)**.

**Fig. S5.**
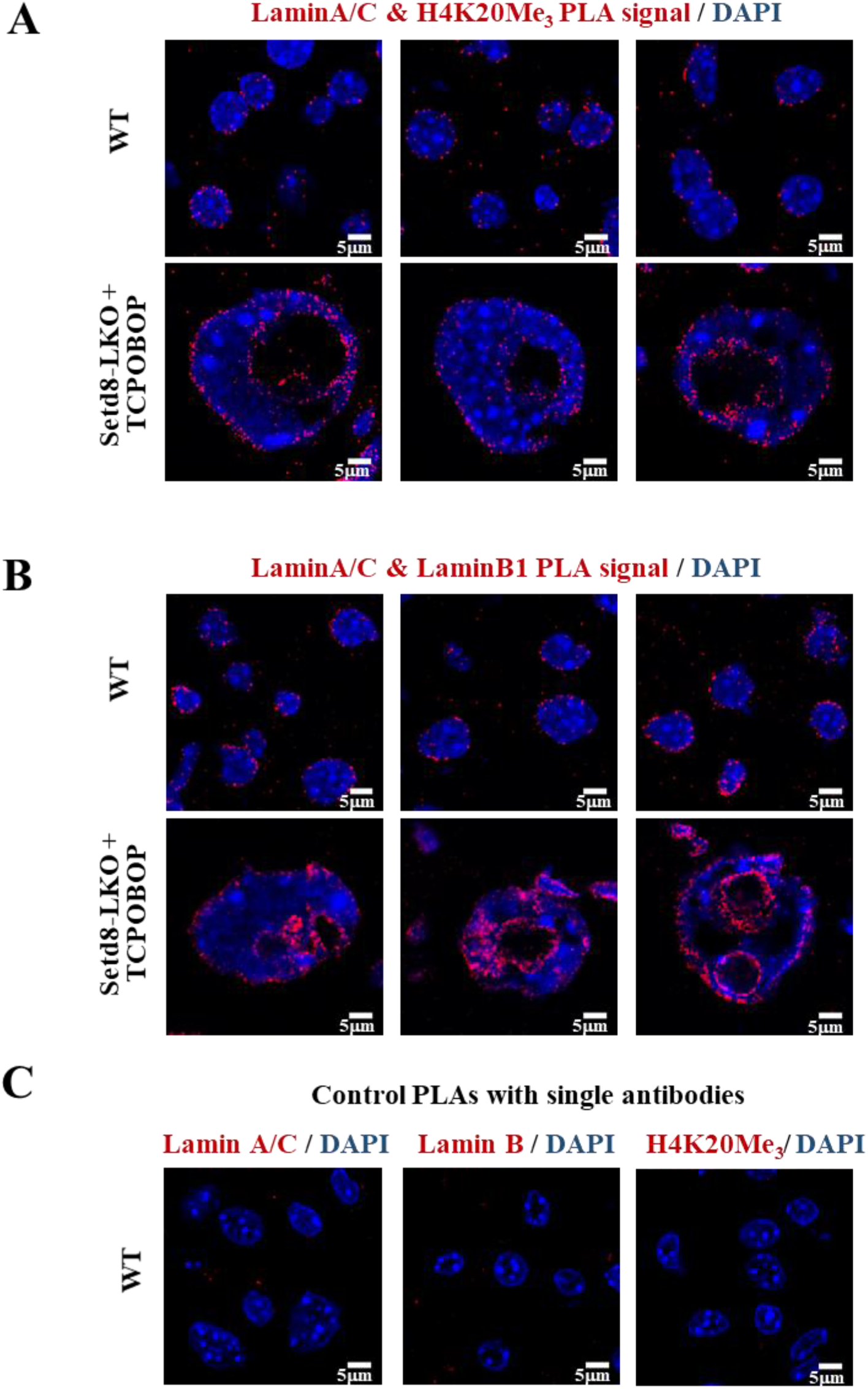
*In situ* detection of H4K20Me_3_-modified nucleosomes associated with nuclear lamina. **(A)** PLA images in liver sections from 3 different wild type and TCPOBOP-treated Setd8-LKO mice using antibodies recognizing H4K20Me_3_ and Lamin A/C. Note the positive PLA signal at both nuclear envelope and internalized vesicle membranes. **(B)** Positive control PLA reaction using Lamin A/C and Lamin B1 antibodies detecting physical proximity of Lamin A/C and Lamin B polymers at the nuclear envelope and at the endonucleotic vesicle membranes. **(C)** Negative control PLA reaction with individual antibodies.

**Fig. S6.**
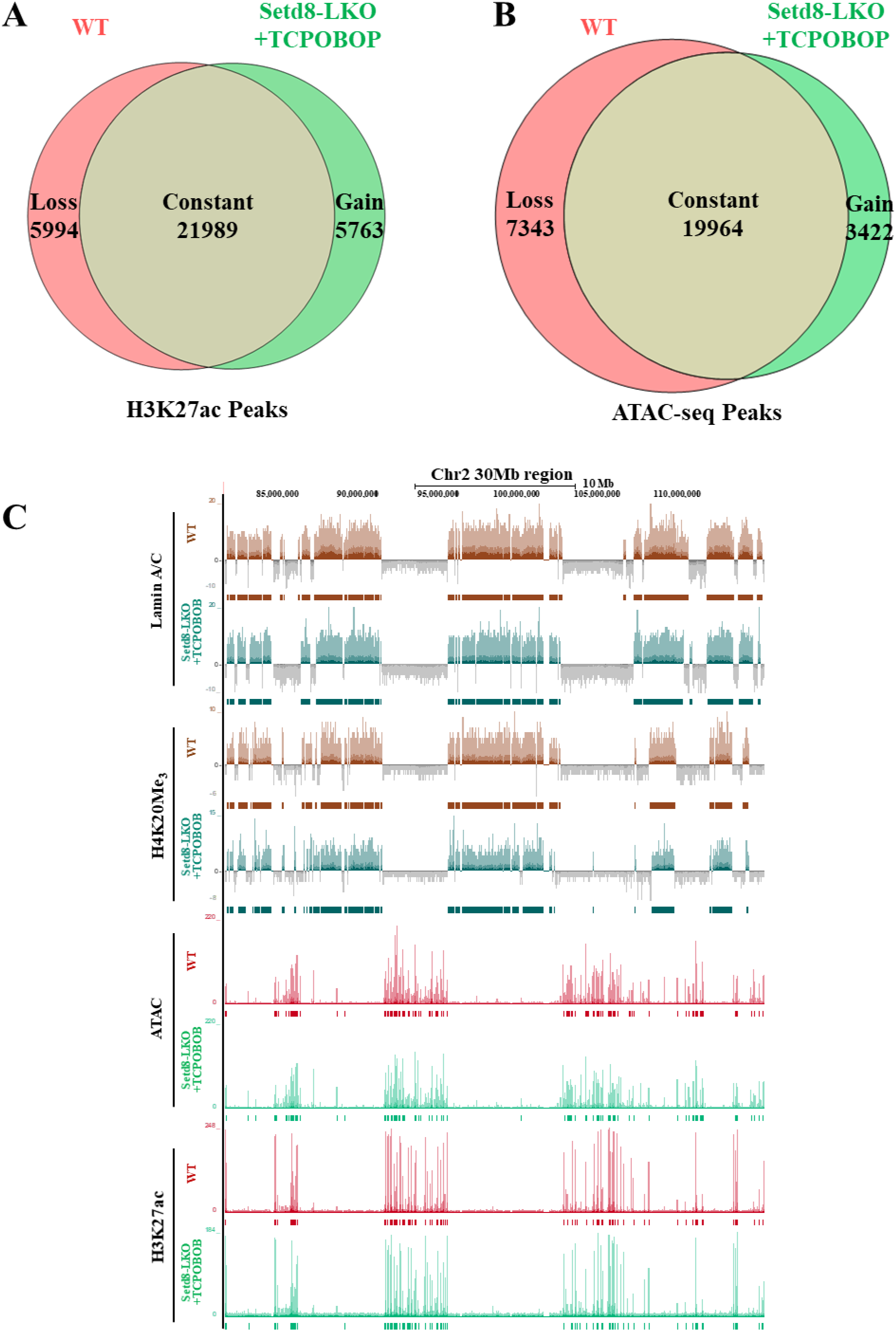
Analysis of transposase accessible, open euchromatin genomic areas in Setd8-LKO hepatocytes. **(A-B)** Venn diagram comparing H3K27ac ChIP-seq peaks **(A)** and ATAC-seq peaks **(B)** in wild type and TCPOBOP-treated Setd8-LKO livers. “Loss” indicates peaks present only in wild type livers, “Gain” indicates new peaks in Setd8-LKO livers. “Constant” correspond to peaks found in both, wild type and Setd8-LKO livers. **(C)** Combined UCSC Genome Browser profile in 30 Mb regions of chromosome 2, showing LADs, H4K20Me_3_-modified heterochromatin domains and euchromatin domains with high density of ATAC-seq and H3K27ac ChIP-seq reads. Note the high-level preservation of the tandemly organized euchromatin and heterochomatin domains in hyperploid Setd8-LKO hepatocytes.

**Fig. S7.**
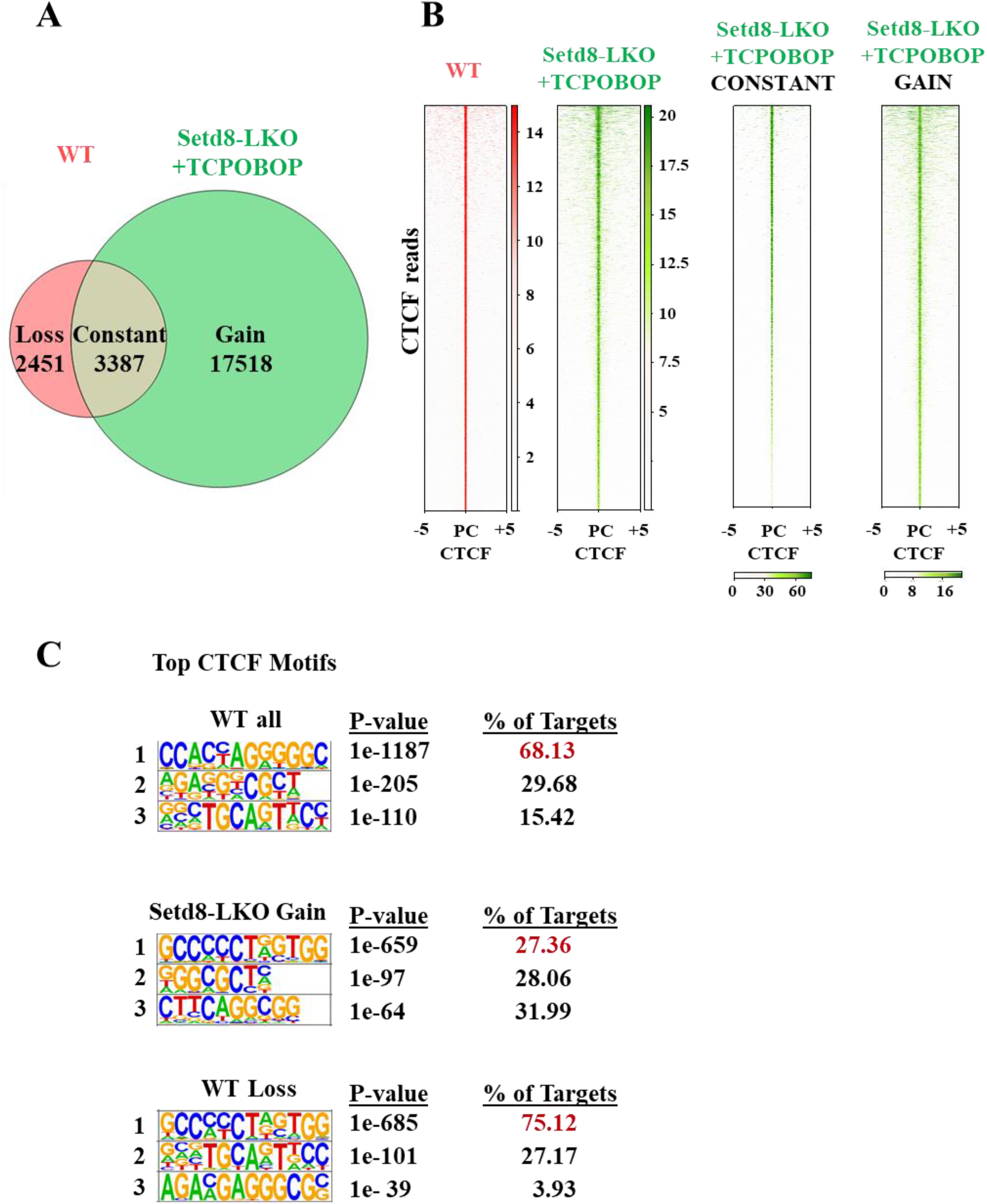
Genome-wide profile of CTCF binding in Setd8-LKO cells. **(A)** Venn diagram showing the overlap of CTCF ChIP-seq peaks between wild type and Setd8-LKO mice 24 hours after TCPOBOP treatment. “Loss” indicates peaks present only in wild type livers, “Gain” indicates new peaks in Setd8-LKO livers, while “Constant” correspond to peaks found in both, wild type and Setd8-LKO livers. **(B)** Binding-intensity heatmaps showing the genomic occupancy distribution of CTCF in 5 kb downstream to 5 kb upstream areas from the CTCF peak center (PC, base pair showing the highest CTCF read pileup) in wild type and TCPOBOP-treated Setd8-LKO livers. The depicted regions were ranked by decreasing CTCF binding strength, as measured by the normalized fold enrichment of reads under CTCF peaks to the respective input regions. Panels at right show analyses restricted to “Constant” peaks and “Gain” peaks. Note the different scales in the heat bars. Note the lower binding strength and the increased number of reads outside the PC of “Gained” peaks. **(C)** Sequence logos of the binding motifs identified under CTCF peaks by de novo motif search. The percent of targets with the canonical CTCF motif (33) are indicated in red color.

**Fig. S8.**
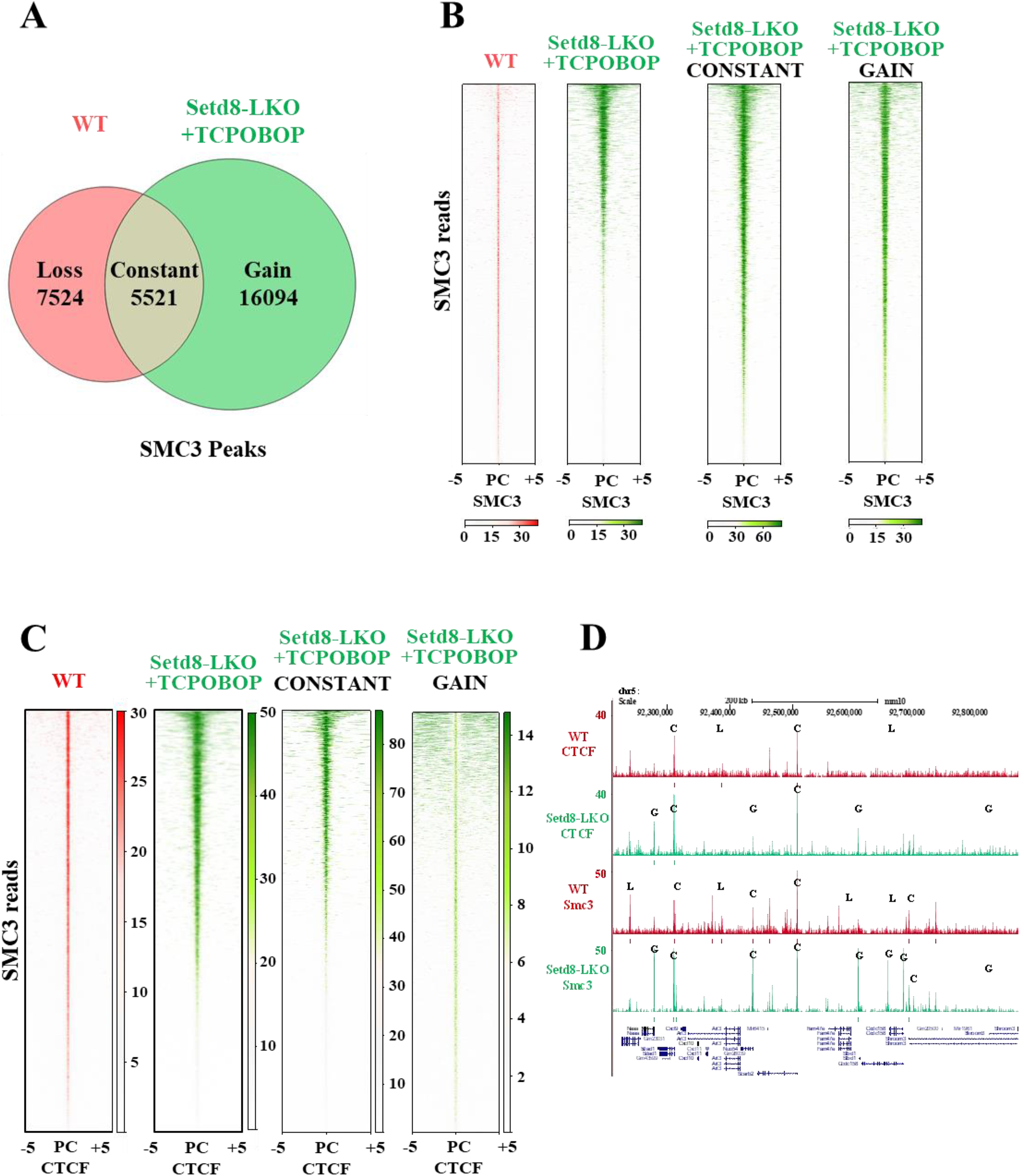
Genome-wide profile of Smc3 binding in Setd8-LKO cells. **(A)** Venn diagram showing the overlap of Smc3 ChIP-seq peaks between wild type and Setd8-LKO mice 24 hours after TCPOBOP treatment. “Loss” indicates peaks present only in wild type livers, “Gain” indicates new peaks in Setd8-LKO livers, while “Constant” correspond to peaks found in both, wild type and Setd8-LKO livers. **(B)** Binding-intensity heatmaps showing the genomic occupancy distribution of Smc3. Heatmaps feature Smc3 reads around called Smc3 peak centers, ranked by decreasing signal strength. **(C)** Binding intensity heatmaps of Smc3 reads in wild type and Setd8-LKO livers relative to the center of CTCF peaks identified in wild type or Setd8-LKO mice. The two panels at right show distribution of “Constant” and “Gain” Smc3 peaks. Note the differences in the color bars indicating relative signal strength. Note the lower binding strength of “Gained” reads compared to “Constant” ones, and their low level accumulation near CTCF locations. **(D)** UCSC Genome Browser profile in 600 kb region of chromosome 5, showing examples for the Loss (L), Gain (G) and Constant (C) categories of CTCF and Smc3 binding profiles in wild type or Setd8-LKO livers.

**Fig. S9.**
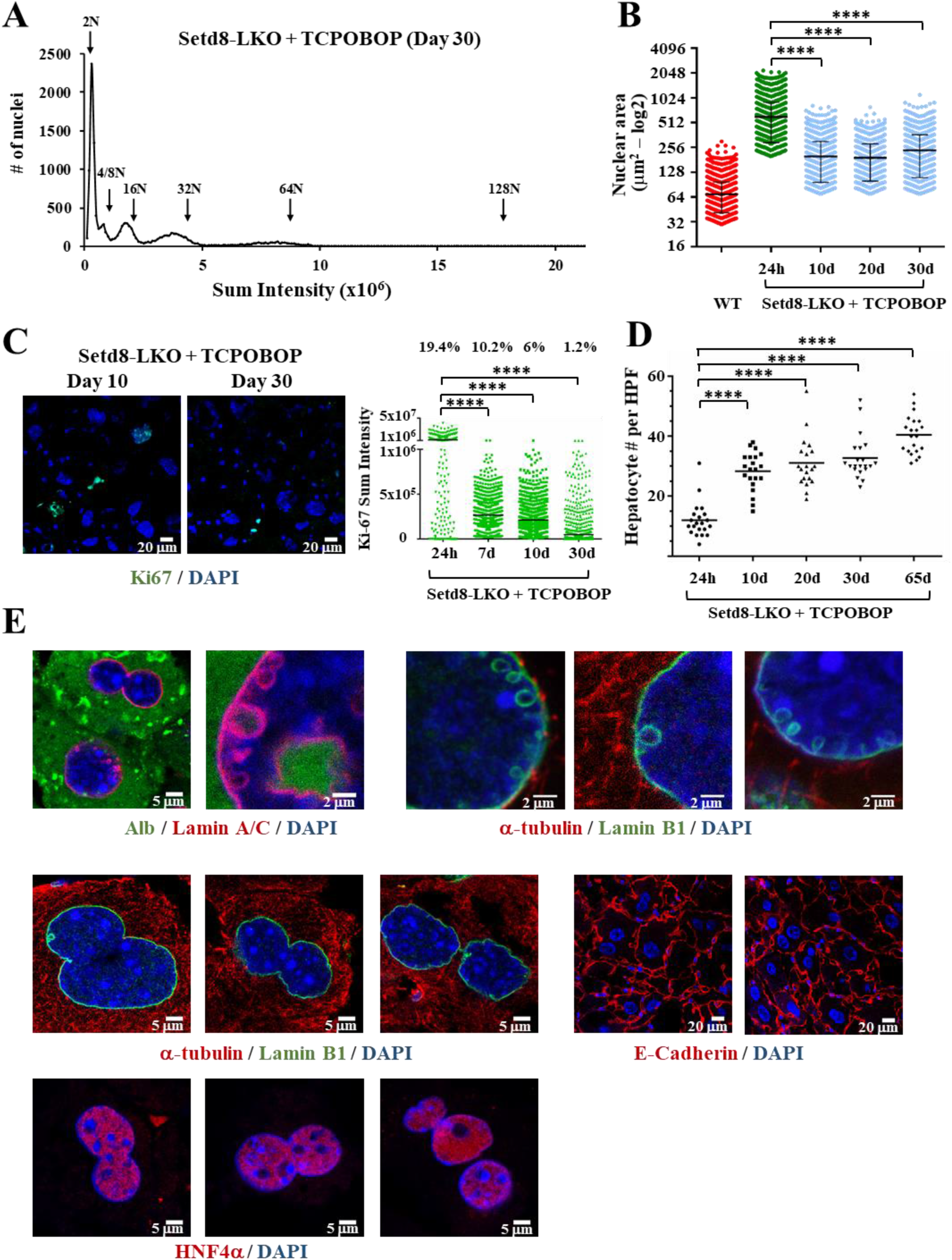
Characteristics of Setd8-LKO livers 20 to 30 days after TCPOBOP treatment. **(A)** Estimation of nuclear DNA content in isolated nuclei from Setd8-LKO livers (n=12569) by propidium iodide (PI) staining 30 days after TCPOBOP treatment. Graphs show high-content-microscopy (HCM) measurements of the PI staining intensity in individual cells. Arrows indicate the chromosomal ploidy of the peaks. The percentage of the cells corresponding to the peak areas were as follows: in 2N=43.2%; ∼4/8N= 8.2%; ∼16N=17.7%; ∼32N=19.2%; ∼64N=8%. Note the reappearance of cells with 2N DNA content and the substantial deviation of the peak centers from locations corresponding to sum intensities with doubling chromosome numbers, which is indicative of aneuploidy. **(B)** Representative images of liver sections stained for Ki67 in Setd8-LKO mice, 10 and 30 days after TCPOBOP treatment. The graph at right shows the quantification of the fluorescence sum intensities in cells stained positively for Ki67 at the time points 24 hours (n=1228), 7 days (n=1593), 10 days (n=1534) and 30 days (n=1495) after TCPOBOP treatment. Note the gradual decrease of the percentage of cells with Ki67 signal above threshold and >5-fold decrease of the median Ki67 pixel intensity in the 7-10-30 days’ time points. **(C)** Comparison of the relative nuclear areas at different times following TCPOBOP treatment. The data for wild type and the 24-hour time point of Setd8-LKO mice are the same as in Fig. S1C, corresponding to the volumes of n=4295 nuclei and n=4095 nuclei, respectively. The data shown in 10 days, 20 days and 30 days after TCPOBOP treatment are measurements from n=3736, n=3538 and n=3690 nuclei, respectively. **(D)** Quantification of hepatocyte numbers in wild type and TCPOBOP-treated Setd8-LKO livers. The graph shows hepatocyte numbers in 20 High Power Fields (HPFs) in Setd8- LKO livers at the indicated time points after TCPOBOP treatment. **(E)** Representative images of Setd8-LKO hepatocytes 20 days after TCPOBOP treatment stained for the cytoplasmic proteins Albumin and α-tubulin, the nuclear protein HNF4α and the cytoplasmic membrane marker E-Cadherin. Note that Albumin and α-tubulin are absent from the small vesicles. Albumin is detected only in some remaining large vesicles. Segregating nuclei have diffuse HNF4 distribution and lack tubulin stained patterns characteristic to mitotic chromatin, which may partition in the same cell (binuclear cells) or separate cells (mononuclear cells). ****p-value<0.0001.

**Fig. S10.**
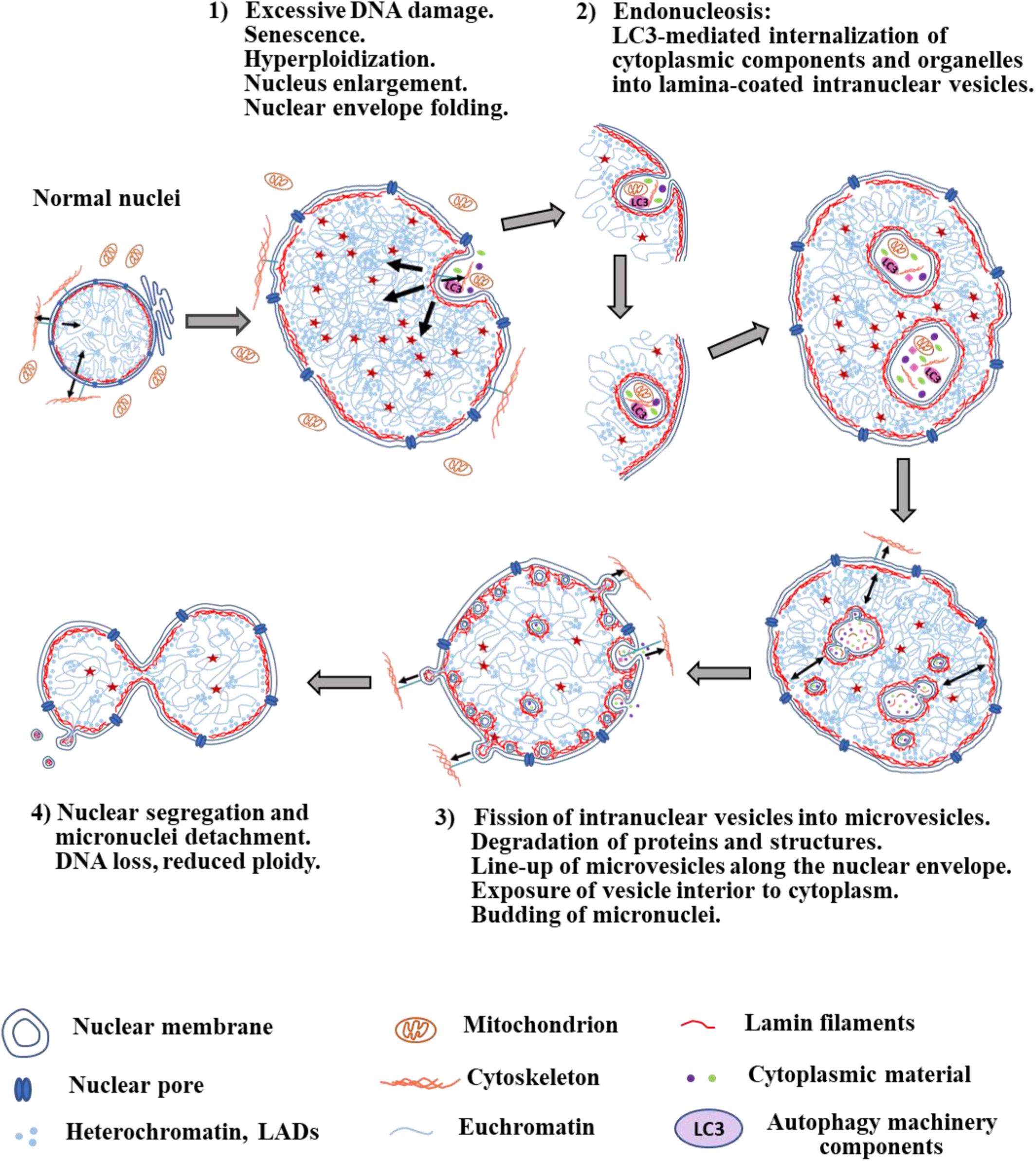
Schematic presentation of the novel senescence features described in this study.

